# Latent Diffusion for Conditional Generation of Molecules

**DOI:** 10.1101/2024.08.22.609169

**Authors:** Benjamin Kaufman, Edward C. Williams, Ryan Pederson, Carl Underkoffler, Zahid Panjwani, Miles Wang-Henderson, Narbe Mardirossian, Matthew H. Katcher, Zack Strater, Jean-Marc Grandjean, Bryan Lee, John Parkhill

## Abstract

Designing a small molecule therapeutic is a challenging multi-parameter optimization problem. Key properties, such as potency, selectivity, bioavailability, and safety must be jointly optimized to deliver an effective clinical candidate. We present COATI-LDM, a novel application of latent diffusion models to the conditional generation of property-optimized, drug-like small molecules. Diffusive generation of latent molecular encodings, rather than direct diffusive generation of molecular structures, offers an appealing way to handle the small and mismatched datasets that are common for molecular properties. We benchmark various diffusion guidance schemes and sampling methods against a pre-trained autoregressive transformer and genetic algorithms to evaluate control over potency, expert preference, and various physicochemical properties. We show that conditional diffusion allows control over the properties of generated molecules, with practical and performance advantages over competing methods. We also apply the recently introduced idea of particle guidance to enhance sample diversity. We prospectively survey a panel of medicinal chemists and determine that we can conditionally generate molecules that align with their preferences via a learned preference score. Finally, we present a partial diffusion method for the local optimization of molecular properties starting from a seed molecule. Conditional generation of small molecules using latent diffusion models on molecular encodings provides a highly practical and flexible alternative to prior molecular generation schemes.

## 1 Introduction

Robust generation of molecules subject to design constraints (i.e., inverse molecular design) has great promise to accelerate pharmaceutical and materials research. Models that generate novel chemical structures with desired properties are a subject of furious development, with methodologies such as genetic algorithms applied to molecular graph structures [Kwon et al., 2021], fine-tuning of autoregressive generators via reinforcement learning [Blaschke et al., 2020, You et al., 2019], sampling from generative models such as variational autoencoders or generative adversarial networks [Jin et al., 2018a, Bilodeau et al., 2022], and sampling from multimodal transformer-based models [Kaufman et al., 2024, Reidenbach et al., 2023, Liu et al., 2023].

To provide practical utility for drug discovery, generative design algorithms need to be highly controllable, computationally efficient, and able to leverage datasets of varying sizes to produce optimized molecular designs in chemical space both near hits of interest and distinct from well-explored regions. Drug design is a challenging multi-parameter problem, and some critical properties, like pharmacokinetic features determining bioavailability, are significantly more difficult to gather data for than others. In this work, we propose the use of latent diffusion models (LDMs) [Rombach et al., 2022] and large pre-trained molecular encoder-decoders [Kaufman et al., 2024] for generative molecule design. While diffusive methods have been applied in molecular graph generation [Vignac et al., 2022] and 3D conformer generation [Satorras et al., 2021, Schneuing et al., 2023, Pinheiro et al., 2023, Huang et al., 2023, Xu et al., 2023], we abstract out the problem of encoding and decoding molecular structures to a large, pre-trained model. This work uses a pre-trained transformer-based contrastive (2D/3D) encoder-decoder trained on a large (∼250M) unconditioned sample of chemical space, which has learned a fixed-dimensional molecular embedding. This architecture is an extension of the COATI model as defined in [Kaufman et al., 2024]. We refer to this application of latent diffusion models to COATI embeddings as COATI-LDM.

Many properties must be optimized to produce a clinical molecule, and in many cases, limited data makes it a few-shot generation problem. Assays that assess absorption, distribution, metabolism, excretion, and toxicity (ADMET) are often costly and therefore the datasets are smaller. Public sources such as the ADMET datasets contained in the Therapeutic Data Commons [Huang et al., 2021] or for human feedback on molecule designs [Choung et al., 2023a] only contain a few thousand samples. Analogous to the work in [Sinha et al., 2021], which describes the application of latent diffusion for few-shot image generation, this work applies the same concept to drug-like chemical space, where data limitations are particularly severe and decoders are less standardized. Latent diffusion methods provide a straightforward and computationally efficient path toward applying a variety of physicochemical and activity constraints to generated molecules.

First, we show that latent diffusion methods can effectively capture the embedding distribution of a large dataset of empirical binding affinity measurements against Human Carbonic Anhydrase II (hCAII) [Sly and Hu, 1995]. Utilizing this dataset, we evaluate the ability of several conditional diffusion approaches to match *conditional* property distributions, i.e., the distribution of embedding vectors conditioned on binding affinity or physicochemical descriptors. We additionally evaluate each method’s level of control over molecular properties via the correlation between conditioning values and the resulting properties of the generated molecules.

We then perform a set of practical benchmarks using empirical binding affinity data, first demonstrating that conditional latent diffusion models provide better coverage of hit-like chemical space compared to highly competitive evolutionary algorithm-based approaches. Additionally, we show that this method is competitive with transformer-based conditioning, despite the latter being trained on a much larger dataset. Furthermore, in analogy to image generation or large language models (LLMs), these conditional diffusion models can be aligned to human preferences (in this case, expert medicinal chemists) for producing molecules with high desirability by using a small dataset. We validate this alignment by prospectively surveying a small panel of medicinal chemists.

We also provide a straightforward method for conditionally sampling near a seed molecule which we call “partial diffusion”: partially noising the seed molecules latent representation and then performing conditional denoising. We show that this method can jointly control molecule similarity, potency, and other properties, and obtain state-of-the-art results on a common constrained molecular optimization benchmark. Finally, we show that particle guidance [Corso et al., 2023a] can be used to generate diverse sets of molecules, raising the possibility of prospectively generating “coverage sets” of chemical space without resorting to clustering on pre-generated chemical sets.

Overall, we find conditional latent diffusion to be a versatile tool for molecular generation under various conditions and disparate datasets. A generative method that separates the data required to condition a molecular property from the data required to learn the structure of admissible chemical space is often vitally important because the data for the former is much scarcer than the data for the latter. Latent diffusion is an amenable approach to this problem, and we present the first application of latent diffusion models to conditional optimization of drug-like molecules using modular conditioning models, trained on separate data sources. We combine a separately-trained invertible continuous embedding of chemical space, derived from [Kaufman et al., 2024], with a straightforward application of conditional latent diffusion methods, and produce a highly performant generative molecular design method.

## 2 Related Works

Herein we offer a summary of generative approaches to drug-like molecule generation, but refer to reviews on this rapidly evolving topic for further detail [Bilodeau et al., 2022, Du et al., 2022]. The vast majority of data-driven generative molecular models focus on direct modification of a molecular structure representation in a stepwise fashion. Evolutionary algorithms [Kwon et al., 2021], which long predate data-driven machine learning but often outperform newer methods [Jensen, 2019], iteratively mutate a molecular structure using operators on graphs or SMILES strings. These methods require the manual curation of a library of chemically reasonable mutation and crossover operators and need to query (possibly costly) oracle functions – such as a docking program or other predictive model – at every step of generation. Methods based on reinforcement learning suffer from the same requirements, performing approximate gradient steps to fine-tune a generative model [Zhou et al., 2019, Blaschke et al., 2020] using feedback from an oracle function.

More recent methods for conditional generative design perform local modifications to a molecular structure via a gradient ascent-like process [Fu et al., 2022] or via a diffusion process on graph structures themselves [Liu et al., 2024]. Generative models relying on 3D structural data generate molecules atom-by-atom [Hoogeboom et al., 2022, Schneuing et al., 2023] or by modifying torsion and bond angles [Corso et al., 2023b, Brocidiacono et al., 2023] of a complete molecular structure to produce realistic binding poses, using datasets such as PDBbind [Wang et al., 2004]. Others have used crystal structures of a target protein binding pocket as conditioning information [Schneuing et al., 2023]. In this work, we focus on the diffusive optimization of a continuous latent vector representation derived from a foundation encoding model, which allows the straightforward application of diffusion modeling without many domain-specific changes.

Methods that leverage a latent molecular representation to perform generative optimization have also been explored in the literature. Early attempts at using variational autoencoders (VAEs) to traverse a latent molecular space [Lim et al., 2018, Gómez-Bombarelli et al., 2018] often suffered from low validity (i.e., generating physically implausible molecule structures) and were often trained on small molecular datasets. More recent VAE methods use molecule graph-specific modifications [Jin et al., 2019a] to the encoder and decoder networks to achieve higher validity. Posterior collapse [Wang et al., 2023] remains a challenge with this technique. GeoLDM [Xu et al., 2023] trains an equivariant VAE on small molecule confirmations, and then performs diffusion on geometrically structured latent embeddings with an equivariant score model. However, the geometric structure of the latent space limits the applicability of well-known diffusion guidance approaches and adds additional constraints on the types of allowable guide functions. 3M-Diffusion [Zhu et al., 2024] performs diffusion on the latent embeddings of a VAE trained on molecular graphs, where the latent space has been aligned to textual descriptions of the molecules using contrastive learning. However, the model is designed to condition on text rather than directly on molecular properties.

More recent methods train autoencoding models from commercial catalogs containing hundreds of millions to billions of synthetically available molecules, which are excellent proxies for chemical realism and accessibility. Recurrent neural networks trained to jointly autoencode molecules and properties [Winter et al., 2019, Hoffman et al., 2022a] or multimodal transformer-based models [Liu et al., 2023, Kaufman et al., 2024] produce a latent code representing a molecule structure. Latent optimization algorithms are advantageous as they decouple the problem of representing chemical data from the construction of generative and predictive models.

Statistically significant experimental measures of performance for generative molecular design are costly to produce, which has frustrated fair comparisons between these methods. Simple benchmarks that estimate chemical validity [Brown et al., 2019] or the ability to produce novel molecules are useful toy examples, but are often unrepresentative of realistic drug design processes [Walters, 2023]. In this work, we focus on modeling an experimental dataset of binding affinities (see Section 4), with a large enough size to calculate Frechet distances. We also survey expert medicinal chemists (Section 5.3) to evaluate our ability to align models to their preferences.

## 3 Methods

Latent diffusion [Rombach et al., 2022] supposes that it is advantageous to generatively model a low-dimensional latent manifold rather than a structured high-dimensional one, given a high-performance pre-trained encoder-decoder. Image diffusion models, for example, perform diffusion in a vector space that can be decoded into an image. In the context of molecule diffusion, we require an invertible encoding for molecular structures, which we describe in Section 3.1. We then pair it with diffusion and flow matching models (Section 3.2) to generate molecule structures with optimized properties. All experiments were trained using a distributed cluster of A100 GPUs on the Nvidia DGX-C compute cloud [Nvidia, online]. Development and training of the encoder-decoder architecture on a 250-million molecule dataset (Section 4) took roughly 20,000 GPU hours, while the top-performing FID diffusion models trained on the same dataset took around 4,000 GPU hours. The associated code repository (https://github.com/terraytherapeutics/COATI-LDM) provides further details.

### 3.1 Molecular Encoder-Decoder

In this work, we rely on a large-scale pre-trained encoder-decoder that maps chemical space (in either 2D graph structures or 3D point clouds) to fixed-length latent vector representations. We adapt the general framework of COATI[Kaufman et al., 2024], in which contrastive learning is used to align 2D and 3D molecular encodings with decoding capability, and extend it by employing a chirality-aware 3D encoder based on Allegro [Musaelian et al., 2023], using DirectCLR [Jing et al., 2022] to control contrastive mode collapse, increasing the latent vector size to 512, and expanding the training dataset. More details on our encoder-decoder are available in Appendix B. We note that the general framework of molecular latent diffusion would function with any sufficiently expressive decodeable molecular representation.

The molecular decoder we employ can complete prompts that contain a property condition, such as `[PROP][LOGP][3][SMILES]`…, which would generate a molecule likely to have a logP of 3. This is another possible modality for conditioned generation of molecules, which we compare against in this work. See Table 6 in Appendix B for a complete list of such available properties. We find in Section 5.4 that the transformer’s property control is quite good; however, applying such a model to new properties requires a costly and laborious fine-tuning and vocabulary injection step that is not required in the latent diffusion framework.

### 3.2 Latent Diffusion

Latent diffusion [Rombach et al., 2022] offers a solution to the problem of generating samples in high-dimensional spaces by learning mappings between lower-dimensional embeddings and Gaussian noise. In this work, we focus on 512-dimensional molecular embeddings {**v**_*i*_ *∈* ℝ^512^} and normally distributed noise 𝒩 (**0**^512^, **I**^512^). This converts the high dimension chemical generation problem into a mathematically much tamer problem of learning a *score network*, 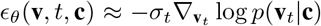, which separates Gaussian noise *ϵ*, from a samples (**v**_*i*_) along a diffusion trajectory (possibly conditioned by conditions, **c**) which maps data samples at time *t* = 0, onto normal uncorrelated samples at *t* = *T* . [Ho et al., 2020] showed such processes can be learned from a straightforward denoising loss,

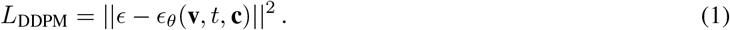

Our results are based on the mathematically convenient formalisms of denoising diffusion probabilistic models (DDPM) and denoising diffusion implicit models (DDIM) [Ho et al., 2020, Song et al., 2022] which are trained with the same loss function.

Flow-Matching (FM) models [Lipman et al., 2023, Ben-Hamu et al., 2024a] generalize the diffusion generative process to a more flexible family of integrated vector fields with adjustable priors and have shown superior performance to other diffusion methods in certain settings [Lipman et al., 2023, Stärk et al., 2023]. We will show some results from optimal transport conditional flow matching, which builds a linear inductive bias into the dynamics mapping data and prior distributions that could lead to more stable training. Besides minor changes to the training process these flow methods can be used to train joint-conditioned generators in exactly the same fashion as DDPM/DDIM, using the same score models as vector fields, and yielding similar performance.

The generative process can be conditioned to produce samples with a desired property in several ways. These techniques were originally developed for use with image classifiers, but are trivially compatible with arbitrary regression objectives. The simplest scheme, joint conditioning, appends variables of interest onto score network inputs during training and sampling. Classifier guidance (CG) [Dhariwal and Nichol, 2021], perturbs the trajectories with a force derived from some desired conditioning classification or regression objective: 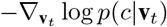 . Better results are obtained by training the regression model or classifier with noised samples and joint knowledge of the point in the noise schedule (*t*) from which a sample is drawn. Throughout this work we use DUE regressors [van Amersfoort et al., 2022] with the same dimension as our latent space whenever we are discussing CG. These are trained using noised samples drawn from the diffusion schedule. The guidance force is: 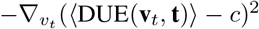. Another possible scheme is classifier-free guidance (CFG) [Ho and Salimans, 2022], which uses the difference between a joint-conditioned and unconditioned score model to guide the diffusion trajectories. D-Flow [Ben-Hamu et al., 2024b] utilizes the deterministic mapping flow matching learns between prior and samples to optimize the condition of a differentiable sample with respect to the initial point on the prior. This makes D-Flow good at achieving strong control while remaining on the data manifold, but the use of a Quasi-Newton solver renders the process much more costly and less predictable because local minima can be found. We experiment with all of these methodologies in the context of molecular latent diffusion, and demonstrate the relative strengths and weaknesses of each method.

Often within a real molecular design project, it is important to generate ‘near’ a previously known molecule due to accessibility of starting materials and increased confidence in structure-activity relationship (SAR) around a known hit. This is another source of strength for latent diffusion over transformer models. Integrating the diffusive generation process forwards and backwards allows a user to explore nearby chemical space while enforcing desired conditions (Fig. 1) and sampling properly on the manifold. Rather than simply adding noise to a latent embedding and decoding, this method traverses the learned manifold of valid molecular structures to find close analogs.

**Figure 1.**
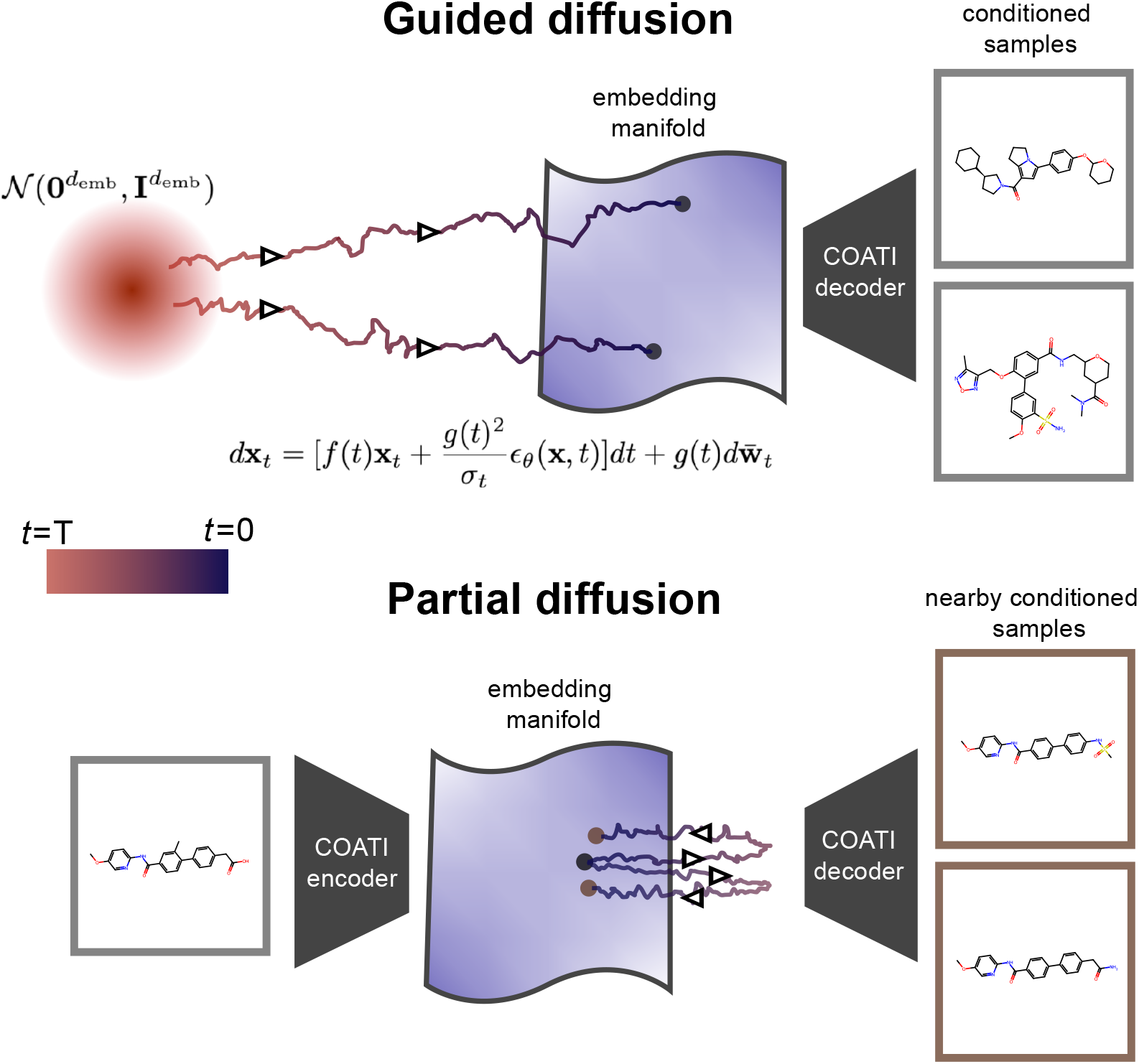
Latent diffusion for molecule generation allows models trained on scarce or non-overlapping datasets to condition generations on a large data manifold which represents a larger and more flexible chemical space. *Partial* diffusion allows one to start with a given molecule and perform a partial diffusion propagation (*t* < *T*) to obtain conditioned samples that are nearby in chemical space.

Score model network architecture has a strong effect on diffusion model performance [Heusel et al., 2017a]. The latent vectors of our encoder/decoder have no translation invariance, and so the U-Net architecture [Ronneberger et al., 2015] usually applied to diffusive image generation isn’t a natural choice. However other components of the architecture are well tuned for diffusion tasks. Given this, we chose to use an architecture in the style of U-Net without convolutions, which is described in Appendix C. The optimization of a score model architecture for this setup is an area for future work.

### 3.3 Preference Alignment

The problem of “aligning” a generative model to human preferences has been widely studied in the context of Large Language Models (LLMs) [Ouyang et al., 2022, Sun et al., 2023, Rafailov et al., 2023] and image generation methods [Wallace et al., 2023, Zhang et al., 2024, Lin et al., 2023]. In the context of small molecular generative models, alignment entails producing molecules that are desirable from a property perspective (i.e., potency) as well as “drug-like”, indicating a high likelihood of oral bioavailability, as well as synthetic accessibility. Medicinal chemists gain a strong intuition for the interplay between these properties. This ‘chemical intuition’ [Kutchukian et al., 2012, Pedreira et al., 2019] allows a medicinal chemist to estimate whether or not a molecule will be worth progressing forward in a program. Historical work into cheminformatics has attempted to quantify this intuition into a series of cutoffs, such as Lipinski’s Rule of 5 or the Quantiative Estimate of Drug-Likeness (QED) [Lipinski, 2004, Bickerton et al., 2012]. More recent work models ‘chemical intuition’ as a preference learning problem, and trains score models on datasets of human-labeled molecule preferences [Takaoka et al., 2003, Sheridan et al., 2014, Choung et al., 2023b].

To align generations to medicinal chemists’ preferences, we adopt the method of [Choung et al., 2023b], which learns a score for a dataset of molecular pairs, where one molecule was picked as “preferable” to the other. Such a model is compact, simple to train, and, most importantly, *improvable* – the scores can easily be fine-tuned by collecting additional preference pairs. This model assumes that pairwise rankings between molecules *m*_*i*_ and *m*_*j*_ is explained by a score function *s*(*m*) that provides an ordering of molecules [Burges et al., 2005]. The learning objective, then, is simply to maximize the likelihood that the chosen molecule has a higher score via a (regularized) binary cross-entropy objective. We fit this model using the latent molecular encoding described in Section 3.1. We then incorporate this information into diffusive generation via simple conditional generation – fitting a smaller conditional diffusion model and using classifier-free guidance, or using a preference score model to condition directly using classifier guidance.

## 4 Datasets

We utilize an ensemble of datasets to probe various properties of latent molecular diffusion models. Outside of Section 5.1, for classifier and classifier-free guidance experiments we use unconditional latent generators trained using an ensemble dataset of over 250 million molecules gathered from publicly available sources of characterized and/or purchasable compounds. This dataset primarily consists of combinatorially enumerated molecule structures using building blocks from Enamine [Grygorenko et al., 2020] as well as publicly available compounds from sources such as ChEMBL [Gaulton et al., 2012], with the objective of providing a close-to-comprehensive survey of practically synthetically accessible chemical space. This dataset was also used to train the encoder-decoder model described above; see Appendix B for more details on its construction.

To examine the ability of latent diffusion to satisfy realistic chemical objectives we use 500K samples of experimental binding affinity data of molecules to a target, hCAII [Sly and Hu, 1995], gathered from a high-throughput binding affinity assay. We also calculate physicochemical properties for these molecules using the RDKit [RDKit, online] to provide examples of conditioning on molecular properties. This dataset is primarily utilized in Section 5.1.

The preference alignment experiments in Section 5.3 use a small dataset (5K molecule pairs, 10K unique molecules) of human-labeled data. In that work, a rating panel of experienced medicinal chemists was assembled and asked to rank pairs of molecule structures based on their desirability. This dataset was first used to train a latent ranking model using the procedure described in [Choung et al., 2023b], which was then used to label data for conditional diffusion model training. Internally, we formed a panel of 3 practicing medicinal chemistry experts to validate that model, finding good agreement. This panel also produced additional preference data and was used to assess the success of conditional generation methods.

See Appendix A for links to these datasets.

## 5 Results

### 5.1 Sampling and Conditional Generation

Generative models are designed to provide an accurate model of their training data distribution. However, in the general case, defining the training data distribution (especially in high-dimensional or discrete cases) is difficult. In the case of generative molecular models, we want to evaluate how well the set of generated molecules represents the distribution of molecules used to train it. A molecular structure, in this setting, is a discrete graph object, computing a parametric distribution of which is difficult. Image generation models are typically evaluated via a comparison between the distributions of low-dimensional embedded samples and training or validation data embeddings. These metrics, such as Frechet Inception Distance (FID) and CLIP Maximum Mean Discrepancy (CMMD) [Heusel et al., 2017a, Jayasumana et al., 2023], compute statistics of lower-dimensional vectors and compare to a reference set. We adopt an analogous approach for evaluating latent molecular diffusion - evaluating coverage using the Frechet distance (FD) of the molecular embeddings generated by the encoder-decoder model described in Section 3.1. While forms of FD are widely used for measuring how generative models fit the distribution of their data, it is typically used in a comparative fashion, as a “good” or “bad” value can be embedding dependent. To contextualise, the current SOTA unconditional FD for image diffusion on the CIFAR-10 dataset (measured using Inception distance) is 1.78 [Kim et al., 2023], but has been progressively decreased from a value of 24.8 from its introduction in [Heusel et al., 2017b]. We use these metrics to compare the relative performance of 3 different diffusion-based conditional generation methods – conditional diffusion, classifier-free guidance, and classifier guidance [Ho and Salimans, 2022, Dhariwal and Nichol, 2021] and additionally evaluate a conditional flow matching model and another flow based method D-Flow [Ben-Hamu et al., 2024a]. We report these distances on the a held-out partition of the hCAII dataset (described in Section 4) in Figure 2. To evaluate the conditional FD, samples are drawn from the model matching the conditions of the samples in the holdout partition.

**Figure 2.**
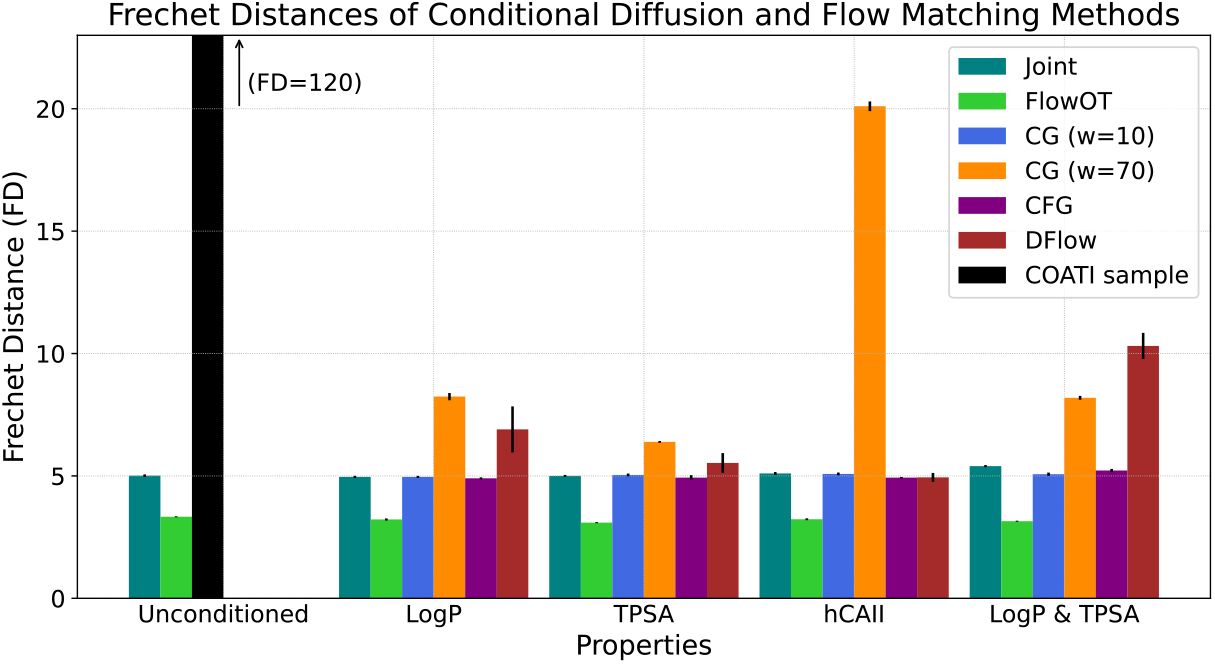
Average FD of different diffusion and flow approaches. Frechet Distances are measured against a 20,000 sample holdout set. The means and standard deviations are computed over five runs of 20,000 samples for each entry, with the exception of the direct sample from COATI which was just run once. *w* refers to the classifier guidance weight used during generation. For each set of the conditions, the conditions of the samples are matched to those in the holdout set. Optimal Transport Flow Matching outperforms all diffusion-based methods, indicating that the learned vector fields induce a better approximation of the training distribution for this dataset.

Separately, we evaluate conditional models’ abilities to produce molecules with desired conditions. We produce a uniform sample of 4000 possible condition values (using the empirical conditional distribution function of the training set values), sample corresponding molecules, and compute Pearson correlations with the desired property. For every condition but hCAII potency, the molecules are decoded into SMILES strings and the properties are computed directly using the RDKit [RDKit, online]. To evaluate hCAII potency conditioning, we predict the value with an MLP trained on the entire hCAII dataset.

We report Pearson correlations between target conditions and the measured properties of the corresponding generations in Figure 3. This and 2 give insights into the generative perfomance of these approaches. We observe that the FDs are fairly low, indicating that they effectively capture the data distribution on this validation set. The three diffusion methods have roughly equivalent FDs across all properties, and the flow matching model performs better across the board. However, we find that better FDs do not lead to better property obedience, with the joint and CFG models having comparable or better correlations on all properties. Notably, CG does not perform as well on property obedience at this weight (*w* = 10), but improved at higher weights. We anticipate that future work will improve upon these metrics, which serve as a baseline for additional developments in generative molecule design on practical datasets.

**Figure 3:**
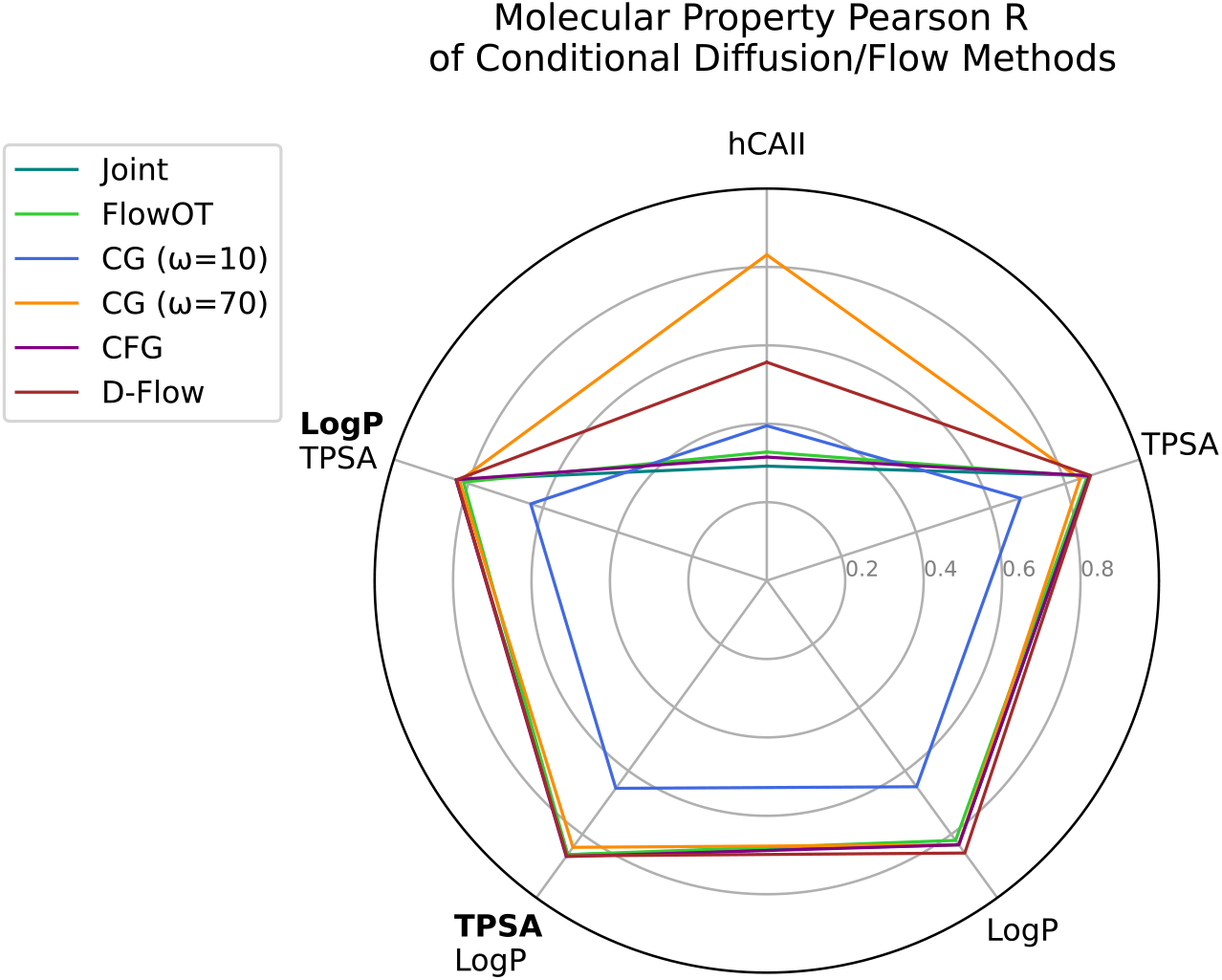
Visualization of Molecule Property/Target Condition Correlations. Values are computed against 4000 condition values that were uniformly sampled over the CDF of the properties, with the exception of the multi-property case, where the samples are treated as independent. For the multi-property case the property the metric is measured on is bolded. We observe that conditioning on more than one property does not degrade the level of control on individual properties. There is limited discrepancy on almost all properties outside of low weight CG. The one exception is hCAII binding, where the classifier guided approaches outperform.

To further explore this we investigate the impact that changing the weight parameters of the guidance methods (CFG and CG) has on FD and condition obedience. While there is no obvious trend when varying classifier-free guidance weights (see Appendix Figure 12) there is a clear trade-off between FD and MAE for classifier guidance, as shown in Figure 4. This type of trade-off between sample fidelity and condition is well suited to a user-adjustable strength of guidance.

**Figure 4.**
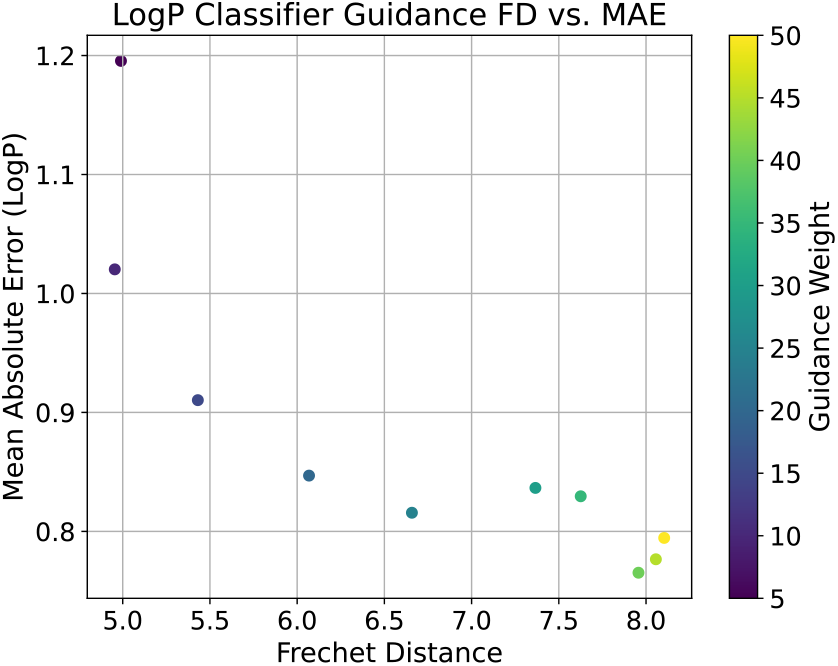
Mean Absolute Error and Frechet Distances of different guidance weights when conditioned on LogP. Increasing guidance weight improves conditioning MAE (i.e., the resulting generations better obey the conditions) but have a higher FD, indicating that the samples are dissimilar to their training set distribution. Frechet Distances are computed with 20,000 samples against the holdout set, while the MAEs are computed on a sample of 4000 conditions uniformly sampled from the LogP CDF of the hCAII training data.

**Figure 5.**
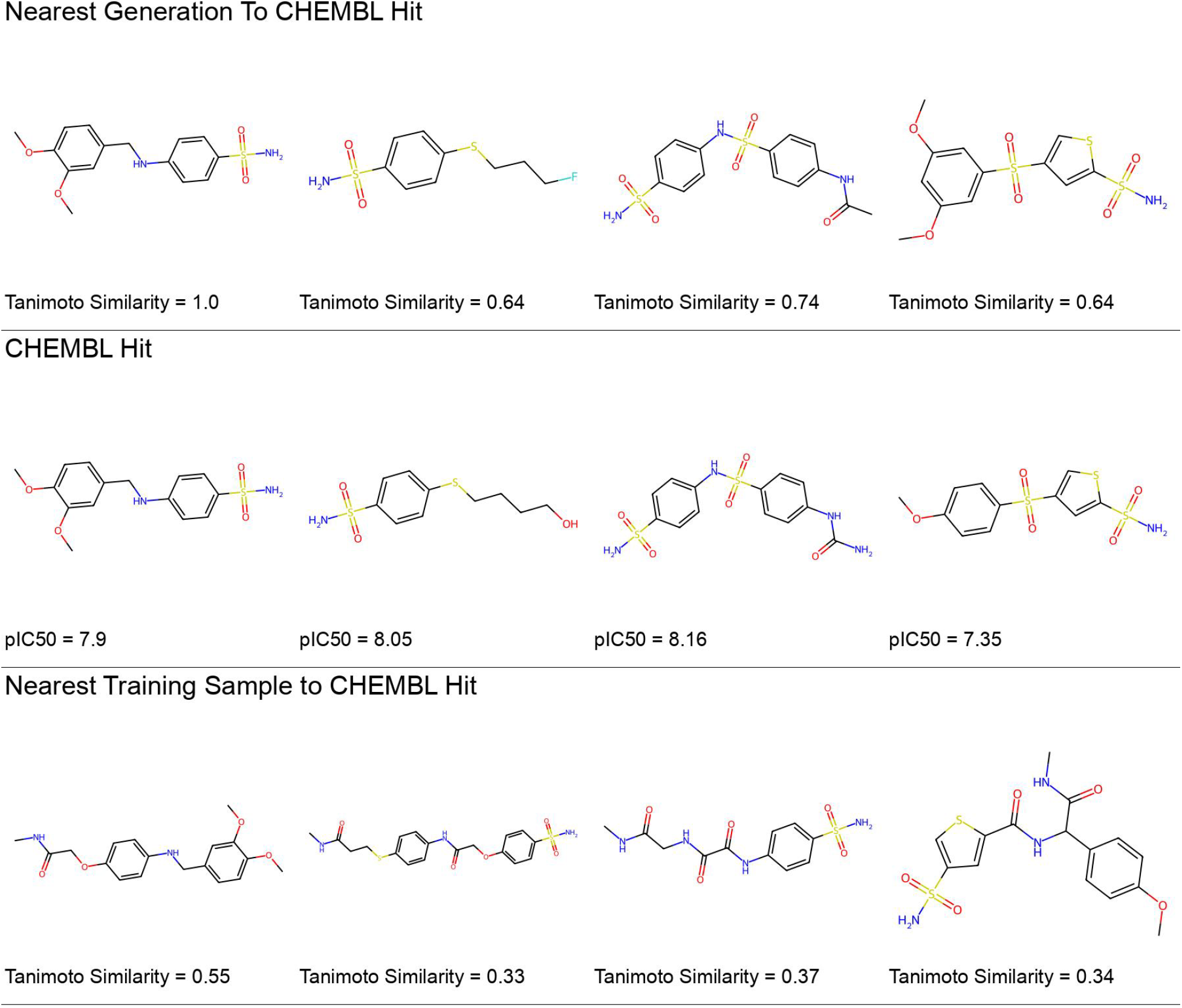
Notable hCAII Generations from the CG with QED model during the genetic comparison experiment. The most similar (ECFP6 Tanimoto) molecule from the public hits dataset is shown for each generation, as well as the most similar molecule from the regressor training data to the public hit.

### 5.2 Comparison to Genetic Algorithms

Genetic algorithms are a strong baseline for molecular generation methods, often outperforming data-driven generative methods on public benchmark tasks [Tripp and Hernández-Lobato, 2023]. Instead of directly modeling the manifold of chemical space, genetic algorithms build up a library of transformations assumed to span molecular space from fragments, and perform a population-based random search in the space of discrete chemical structures to maximize an objective function.

We construct a genetic generative baseline using a pool of fragments furnished by making successive cuts to molecules in ChEMBL 32 [Mendez et al., 2018] and collecting fragments with ≤12 heavy atoms. During each generation, molecules are produced by joining two or more fragments along the scissile atom positions, as determined by a SMARTS match, and the top-scoring percentile is disassembled into an enriched fragment pool for the next generation.

We perform a head-to-head comparison between this fragment-based evolutionary approach and a diffusive generative model to elucidate the strengths and weaknesses of each generative method. To better evaluate the methods in a hit-finding context, generations were assessed by their coverage of a set of held-out “binding hits” from the hCAII data. This set was generated by sorting the top 10% of binding values and putting every other molecule into the holdout set. Additionally, the models were evaluated on 544 public hCAII active compounds (pIC50 ≥4) that were both not in training data and not constrained by the building block chemistry of the training data. Coverage was determined by the proportion of each evaluation set which passed a threshold of Tanimoto similarity [Bajusz et al., 2015] on Morgan ECFP6-2048 fingerprints [Morgan, 1965] to fifty thousand generated molecules for each approach. For example, a coverage of 15% means that 15% of the evaluation set had close analogs produced by the generative method.

We benchmark against three genetic algorithm variants – the first uses only the predicted hCAII binding affinity as the objective function, while the second utilizes a quantitative estimate of drug-likeness (QED) [Bickerton et al., 2012] cutoff (QED ≥0.5) as an extra condition when selecting parents, and the third equally weights predicted hCAII binding affinity and QED in its objective function. After 10 generations where the top 10% of molecules are the parents to further generations, a final selection is made from the top 50K molecules of any generation.

To compare against the genetic methods, which are not constrained to sample within the data distribution, we trained an unconditioned model on the large ensemble dataset (Section 4). We used the unconditional model within the CG and CFG setups, training a hCAII regressor for CG and a hCAII conditional diffuser for CFG. For comparison against the genetic algorithm’s use of QED, one CG variation included a conditional QED diffuser with equal weight to the hCAII diffuser. We note that the “public hits” are in ChEMBL – i.e., 500 out of 1M sources of fragments used to produce the genetic fragment set - which could be seen as an unfair advantage for the genetic approach. However, we believe this comparison is still the most interesting, most closely reflecting industrial practice.

In the domain closest to the hCAII training data - the held out Terray binding affinity data, CFG outperformed CG and genetic approaches (Table 1) by generating more somewhat-similar (Tanimoto ≥0.4) and similar (Tanimoto ≥0.7) compounds. We hypothesize that the conditional diffuser utilized in CFG directly models the data distribution rather than simply providing a gradient signal to guide a diffusive generator. This provides an interesting companion to the results observed in Section 5.1, where there is a clear trade-off between distribution matching and conditioning for CG but no such trade-off for CFG.

**Table 1:** Percent of known-potent held-out molecules with neighbors to 50K generations from diffusion methods and genetic algorithms. We find that the classifier-free guidance-conditioned diffuser is best able to recover hits in the hCAII data, which is unsurprising given that its conditioning model was trained on similar data. Classifier guidance, on the other hand, is best able to recover molecules on a public dataset.

| Partition | Tanimoto Cutoff | hCAII Train | CFG | CG | CG w/ Joint QED | Genetic | Genetic w/ QED Cutoff | Genetic w/ Joint QED |
| --- | --- | --- | --- | --- | --- | --- | --- | --- |
| Binding Hits | 0.5 | 99.6 | **42.7** | 5.6 | 10.9 | 0.9 | 1.4 | 1.5 |
|  | 0.6 | 75.7 | **15.3** | 0.9 | 2.1 | 0.1 | 0.2 | 0.2 |
|  | 0.7 | 28.6 | **4.3** | 0.1 | 0.3 | 0 | 0 | 0 |
| Public Hits | 0.5 | 0.6 | 1.1 | 6.1 | **15.1** | 7.2 | 7.2 | 5.0 |
|  | 0.6 | 0 | 0 | 2.4 | **5.7** | 1.8 | 2.6 | 1.3 |
|  | 0.7 | 0 | 0 | 0.6 | **2.0** | 0 | 0.2 | 0 |

For the small set of non-Terray hCAII hits, CG generations showed better coverage than CFG and genetic approaches for “somewhat-similar” and “similar” Tanimoto cutoffs. Molecules from the public hCAII hits are not confined to building blocks and chemistries of the binding affinity assay. Besides the fact that they are known potent binders of hCAII, there is no reason to expect that 50K samples of a generative model would cover the space of these molecules. Even so, we maintain that coverage of known binders is a useful and quantifiable measure of generative generalization. We observe that the the coverage gap between genetic generation and diffusion narrows on these molecules far outside the classifier’s dataset. Diffusion-based methods still outperformed, with substantially less wall-time.

The stark performance difference between CFG, genetic and CG on the “in-sample” but held out HCAII data vs. public test samples highlights how difficult it is to quantitatively compare molecular generation approaches because statistics do not converge well even with larger-than-average datasets. In practical applications training data is the limiting factor, and methods like CG and genetic algorithms which are less expressive, and therefore less overfit, easily outperform.

### 5.3 Medicinal Chemistry Preference Alignment

Table 2 shows correlations between conditioned preference scores and inferred preference values using conditional diffusion models trained on a RankNet-labeled hCAII dataset. To produce these labels, we used the trained preference model outlined in Section 3.3 to label the hCAII dataset described in Section 4, and trained a conditioned diffusion model on the scores. These were then used to perform classifier-free guidance as in Section 3.2. We observe decent control over preference, indicating that the learned ranking scores are easily incorporated into a diffusion model. We also find that conditioning ability is not significantly degraded when conditioning on preference in addition to another property of interest.

**Table 2:**
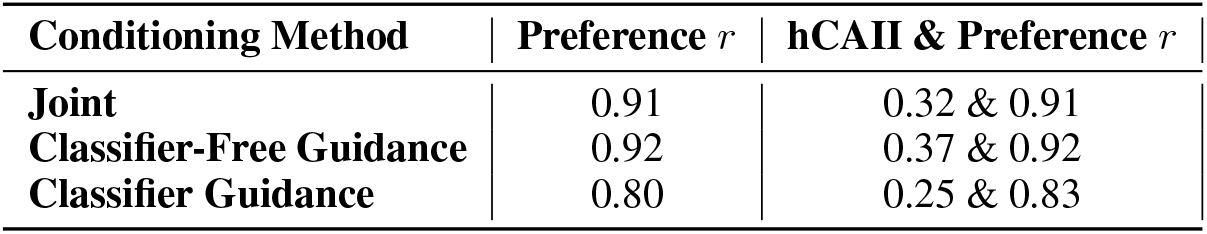
Conditional control over the learned molecule preference score described in Section 3.3. 2048 samples from a uniform distribution were used to condition the diffusion methods, and the resulting vector was used to infer a learned preference score. We see that classifier guidance and direct joint conditioning have better control over preference, and, like with our other properties, conditioning on preference does not degrade conditioning on another property.

| Conditioning Method | Preference $r$ | hCAII & Preference $r$ |
| --- | --- | --- |
| **Joint** | 0.91 | 0.32 & 0.91 |
| **Classifier-Free Guidance** | 0.92 | 0.37 & 0.92 |
| **Classifier Guidance** | 0.80 | 0.25 & 0.83 |

We performed a qualitative analysis of preference conditioning - sampling molecules with high and low desirability as well as jointly conditioning on hCAII potency. Figure 6 contains a t-SNE plot demonstrating that the conditoning produces dissimilar chemical spaces. We see, primarily, that low-desirability molecules appear very large and synthetically complex, with unstable substructures.

**Figure 6.**
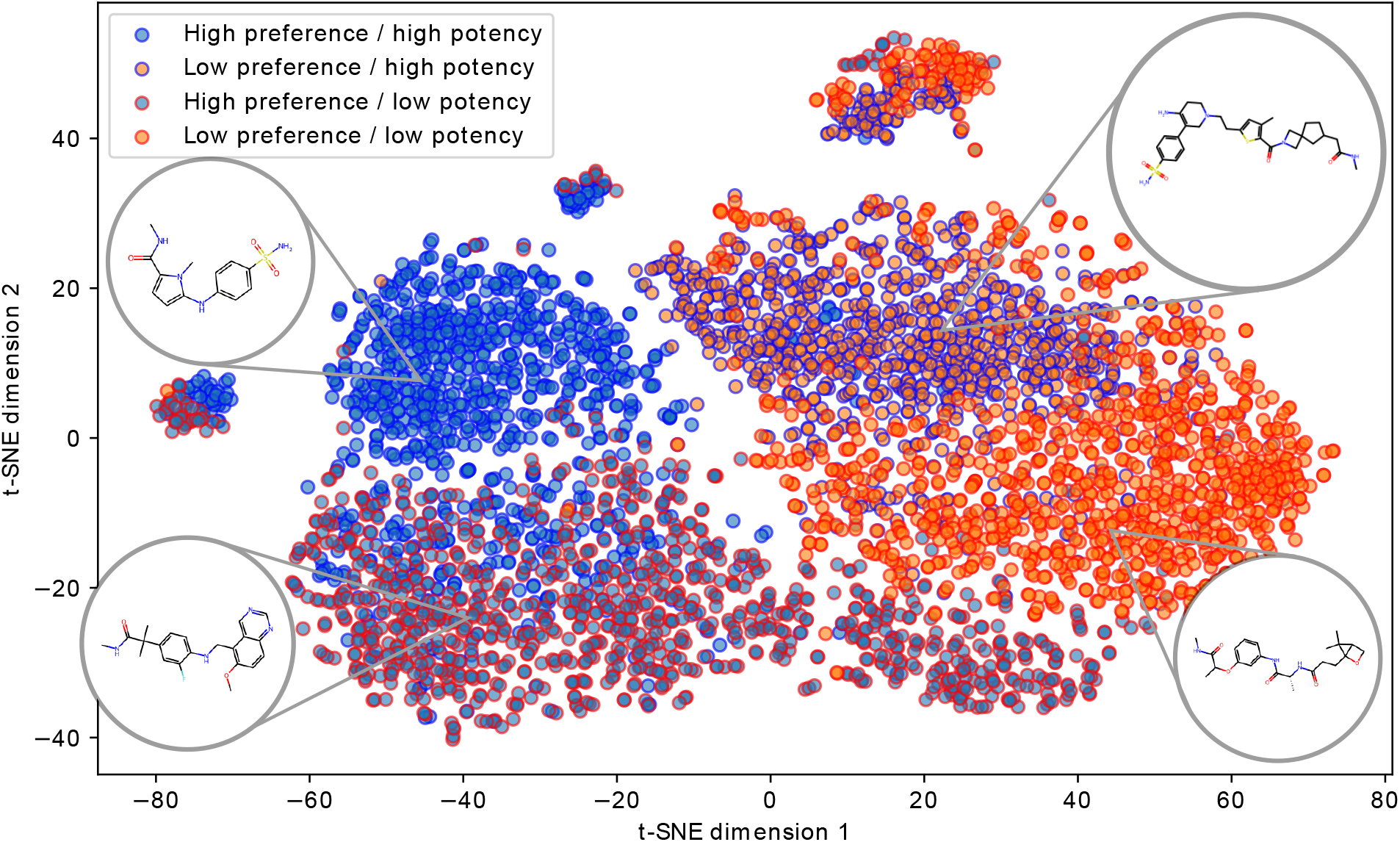
A t-SNE plot of 4096 samples from a CFG process, conditioned jointly on high and low preference as well as high and low hCAII potency. We see that the four conditions produce a fairly disjoint chemical space, with low preference samples appearing larger and more lipophilic, and high potency samples possessing chemical motifs that are present in known binders.

To produce a statistical assessment of our ability to condition on preference, we applied a simplified version of the survey methodology found in [Choung et al., 2023b] to gather data from a group of 3 medicinal chemists. A Streamlit [Snowflake, 2024] app (see Figure 14 for an interface example) presented chemists with a random pair of molecules, who were prompted to select which molecule would be the most desirable. We first performed a survey to confirm that the preference model learned from public data matched the preferences of our medicinal chemistry team. We created a composite dataset consisting of samples from ChEMBL, randomly enumerated combinatorial molecules, and random decodes created by sampling from the molecular decoder described in Section 3.

After we had confirmed that a preference model trained on public data agreed with our teams’ preference (see Figure 13), we designed a subsequent experiment to prospectively test generative preference alignment. Using the low-data conditioned diffusion model described in Section 5.3, we used classifier-free guidance to generate two sets of molecules: (1) a set of molecules with high preference (preference scores 2 standard deviations above the mean) and (2) low preference (preference score between 1 and 2 standard deviations below the mean). We then presented our survey participants with 150 (randomized) pairs using the same methodology as our first-round experiment, without revealing that each pair contained one low-preference and one high-preference molecule. We noticed qualitatively that conditioning for extremely low preference (i.e., producing molecules with the minimum possible preference values) was too obvious – molecules tended to be either macrocylic or very large and lipophilic. To get a fairer survey of preference alignment, we weakened the low-preference condition to better represent drug-like chemical space.

As a simple evaluation of whether or not we can practically enrich generated molecular sets for medicinal chemist-preferred molecules, we then computed the fraction of reported pairs where the high-preference-score molecule was preferred. Across all observations, the high-preference generation was preferred over the low-preference generation in 76.8% of responses, without any fine-tuning using data from our chemistry team. We found that pairs where the chemists disagreed with the conditioning tended to have strange or unstable “high-preference” molecules (See Appendix D, indicating that the preference model can benefit from additional feedback data.

An off-the-shelf preference dataset, alongside this conditioning framework, allows us to produce molecules that are more in line with medicinal chemists’ preference with a minimal amount of tuning. In future work, we anticipate incorporating active feedback from medicinal chemists to improve the quality of generated molecular structures.

### 5.4 Comparison to Transformer Conditioning

The frozen molecular encoder-decoder used in our latent diffusion scheme was trained to jointly encode and decode several molecular properties (the full list of properties can be found in Appendix B) and can be used as a tool for conditioned generation. Latent diffusion is still extremely practical, because of its low data demands, more efficient training, and incremental improvability. Still, the generation quality between these methods merits comparison.

In Figure 7, we compare the distributions of property-conditioned generations for the following properties: topological polar surface area (TPSA) and the fraction of C atoms that are sp3 hybridized (fsp3). The values at which each property was conditioned are given in the x-axis and we generate 256 samples with each method at each value. In Table 3, we report the overall root mean squared error (RMSE) across these specific property conditions, where the multi-property conditioning (TPSA & fsp3) considers all such property value combinations and the RMSE is evaluated separately for each property. The total training set size is also given for context, where the COATI autoregressive generation model was trained on over 500x more molecules than the diffusion models. Despite the much more limited chemical training space, we find that diffusion-based models can sometimes outperform the corresponding autoregressive generator on these chemical properties. Additionally, for the COATI autoregressive generator, we find that performance degrades for the multi-property conditioned generations, whereas in the diffusion models, the performance is better maintained.

**Figure 7.**
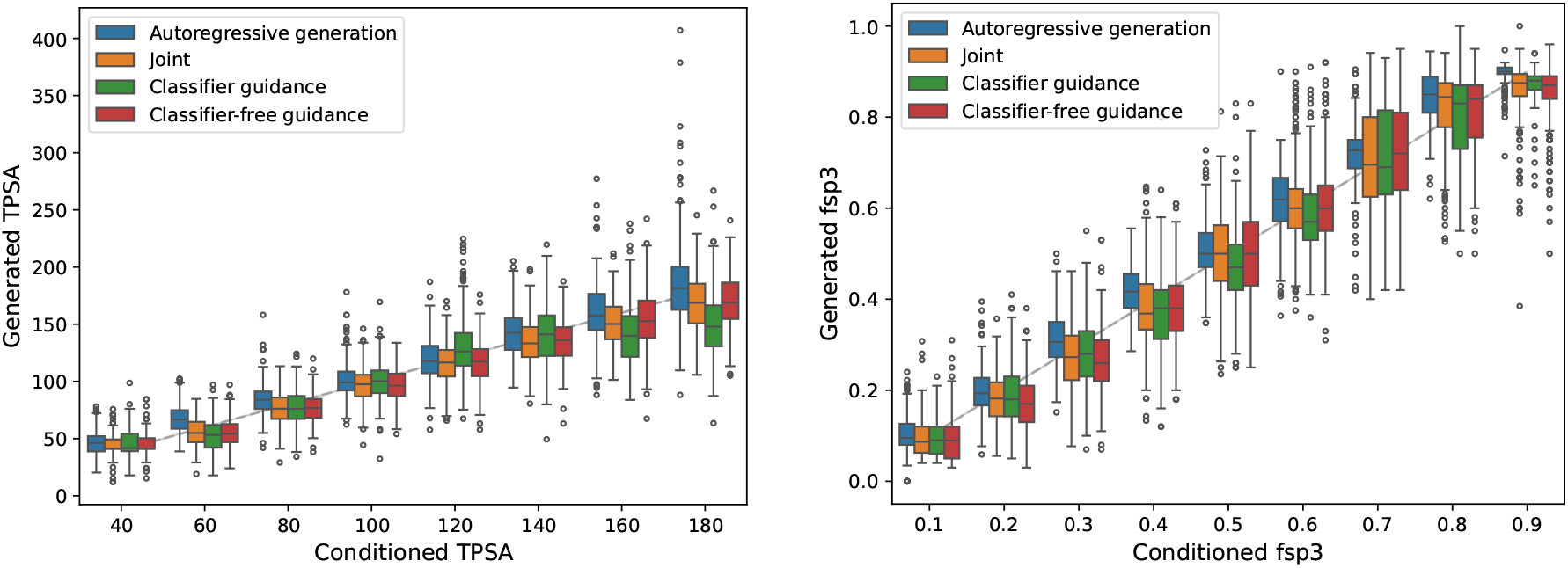
Box and whisker plots (with outliers marked as circles) of generated molecule properties versus the conditioned value for fsp3 and TPSA. Each distribution is over 256 condition-generated samples. The dashed gray line represents a perfect agreement. RMSEs are given in Table 3. Both methods perform well here, although the difficulty of retraining a large autoregressive model on new properties obviously precludes its utility for arbitrary generative tasks.

**Table 3:**
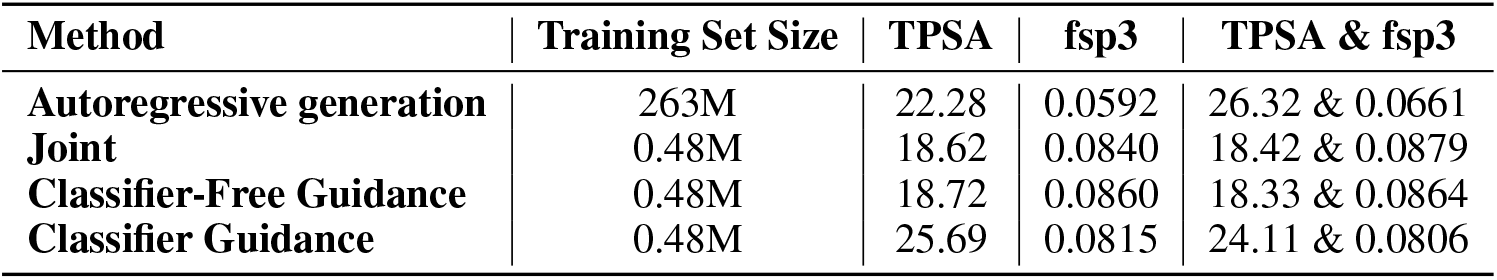
RMSE of conditioned properties in latent diffusion models vs. the parent transformer. We see that diffusion models perform comparably to prompted autoregressive generation with a fraction of the training data.

### 5.5 Nearby Sampling

Small molecule drug design is typically pursued as an iterative cycle of small modifications to an existing “lead” compound. The data domain of predictive molecular models is highly dependent on the chemical space used to fit them, so generating new, in-domain molecules with property improvements is highly relevant to progressing a drug development program. Typically, these modifications are made in attempts to modify suboptimal properties of the lead compound - e.g., improving activity or solubility, or removing toxicophores without reducing activity.

We approach this task as a partial diffusion process – adding limited noise to a starting point using the diffusive forward process, and then using the conditional score model to denoise back into the structured latent space. The number of noise steps, as shown in Figure 8, allows control over the resulting molecules’ similarity to the start point. Classifier-free guidance performed poorly on LogP optimization, failing to effectively improve LogP while maintaining similarity to the starting point. Classifier guidance better achieved this desired behavior. hCAII binding affinity follows a similar pattern. An example of varying the number of forward diffusion steps can be found in Figure 9. This toy example shows the essential trade-off between adding noise and modifying properties - adding additional noise allows for optimization of the property in question (in this case, lipophilicity), but traverses further away from the seed molecule.

**Figure 8.**
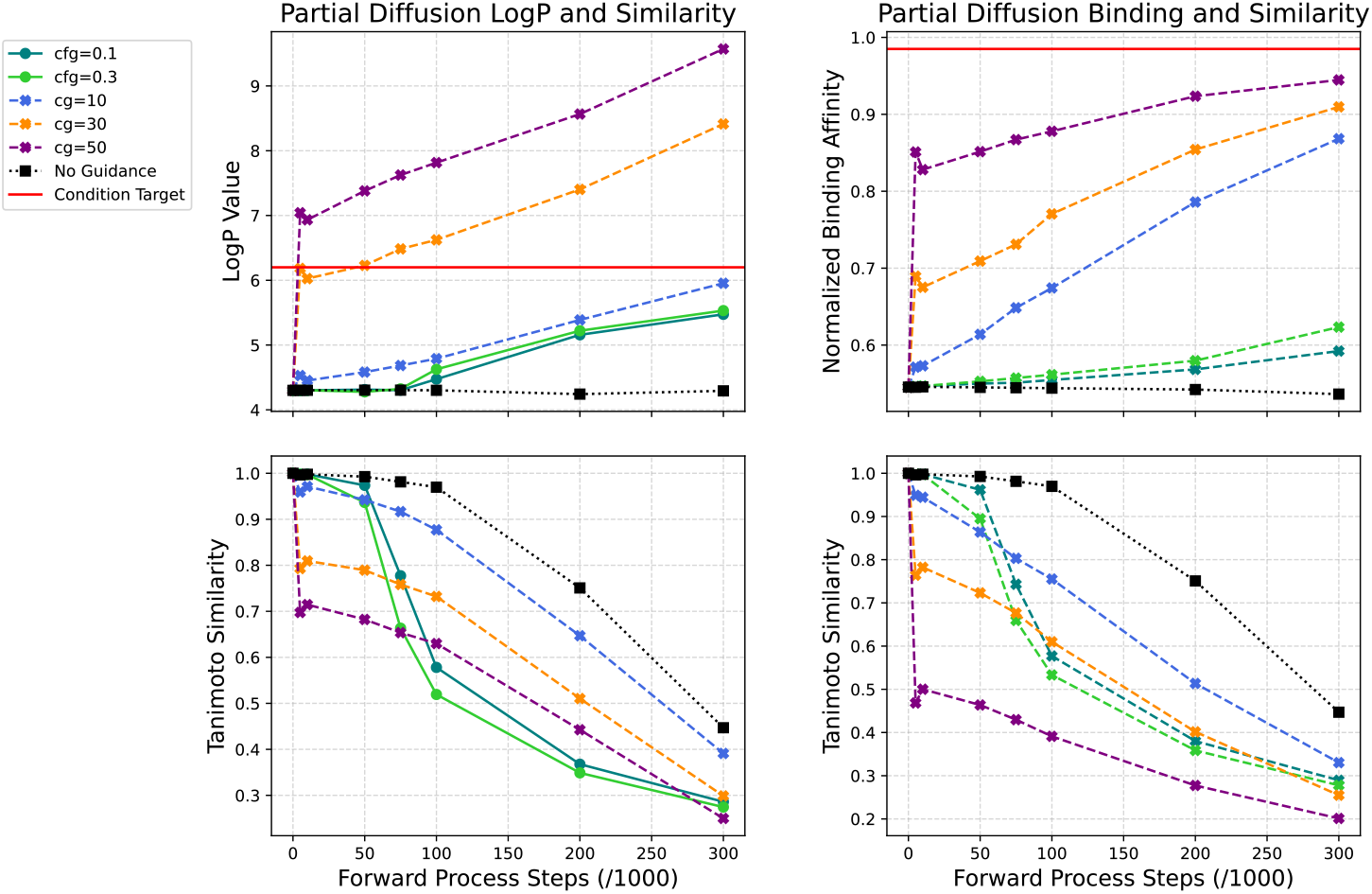
One use case of partial diffusion is to control the similarity of generations relative to a starting point, while also optimizing a desired property. We show this for the maximization of LogP and hCAII binding affinity. These plots show the average Tanimoto Similarity of 1000 diffusion trajectories to their randomly sampled starting molecules and the average property values of the molecules generated by the trajectories. Increasing the number of reverse diffusion steps reduces the similarity to the starting molecule while increasing the conditioned value. Lower weightings of classifier guidance offer a reasonable trade-off of property optimization and similarity to starting point. Classifier free guidance is less effective in this framework.

**Figure 9.**
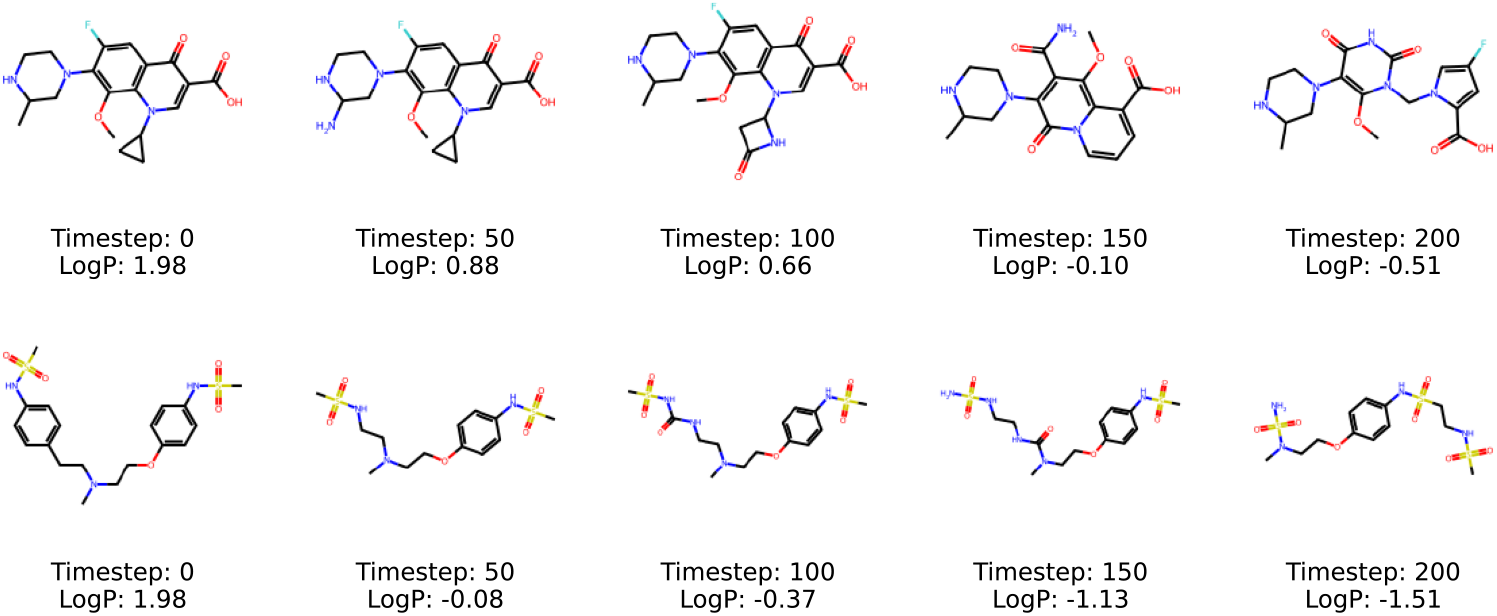
Varying the number of reverse diffusion steps and performing reverse diffusion with classifier guidance allows control over how similar output molecules are to a starting point, trading off property optimization with similarity. Here are examples of molecules generated by partial diffusion on two FDA approved drugs gatifloxacin and dofetilide. Classifier guidance was used to reduce LogP.

This setup was evaluated on a generative benchmark from [Jin et al., 2019b] where, given a set of starting molecules with QED ∈ [0.6, 0.8], the goal is to generate a molecule with QED ≥ 0.9 with a Morgan fingerprint Tanimoto similarity to the starting molecule ≥0.4. Our methodology for the benchmark is detailed in appendix E. As shown in Table 4, latent molecule partial diffusion with COATI-LDM achieves state-of-the-art performance. Notably, this is achieved without directing the process with Tanimoto similarity at all, which differs from MolMIM and QMO which both incorporate it into gradient-free optimization routines, which is a handicap for the diffusion approach. In partial latent diffusion, the relative distance on the latent manifold resembles a Tanimoto similarity, and restricting the number of noise time steps will naturally tend to limit the corresponding Tanimoto similarity deviation of resulting generated molecules.

**Table 4:** Success percentages for the conditional nearby generation task defined in [Jin et al., 2019b]. Given a set of starting molecules with QED ∈ [0.6, 0.8], the objective is to generate a molecule with QED ≥0.9 with a Morgan fingerprint Tanimoto similarity to the starting molecule ≥0.4. Partial diffusion using COATI vectors achieves state-of-the-art performance.

| Method | Success (%) |
| --- | --- |
| JT-VAE<sup>[Jin et al., 2018a]</sup> | 8.8 |
| GCPN<sup>[You et al., 2018]</sup> | 9.4 |
| MMPA<sup>[Dalke et al., 2018]</sup> | 32.9 |
| VSeq2Seq<sup>[Bahdanau et al., 2015]</sup> | 58.5 |
| VJTN+GAN<sup>[Jin et al., 2018b]</sup> | 60.6 |
| AtomG2G<sup>[Jin et al., 2020]</sup> | 73.6 |
| HierG2G<sup>[Jin et al., 2020]</sup> | 76.9 |
| DESMILES<sup>[Maragakis et al., 2020]</sup> | 77.8 |
| QMO<sup>[Hoffman et al., 2022b]</sup> | 92.8 |
| MolMIM<sup>[Reidenbach et al., 2023]</sup> | 94.6 |
| **COATI-LDM** | **95.6** |

### 5.6 Diverse Generations via Particle Guidance

Encouraging the generation of *diverse* molecular sets is an important task in molecular design, in particular during the early stages of drug discovery when lead series have not been solidified. Users may want to avoid the intellectual property space of a known drug, or rapidly assess SAR around a scaffold with as few measurements as possible. Practitioners of cheminformatics have used a method called Directed Sphere Exclusion [Gobbi and Lee, 2003] for decades to diversity filter ensembles of molecules. This method defines a sphere around a seed compound (typically via a Tanimoto distance radius), and assigns all molecules within that sphere to that compound’s cluster. This procedure occurs iteratively until all molecules are clustered, producing a set with enhanced pairwise Tanimoto distances.

Latent diffusion also provides a convenient mechanism to build diversity into generated molecular samples. We employ the concept of particle guidance [Corso et al., 2023a] which simply adds a repulsive guidance force between samples at each timestep while sampling. Figure 10 is a t-SNE plot of 256 generations at varying levels of particle guidance sampled from a jointly conditioned DDPM model for hCAII potency and LogP values between 1.3 and 1.8. The plot depicts training samples, generated samples, and hits from the hCAII interleaved holdout set. Several known potent held-out molecules, especially those that are “isolated” from other hits, are only covered nearby with a sample when particle guidance is applied. These are shown in the figure with the t-SNE plotted points encompassed by dashed circles.

**Figure 10.**
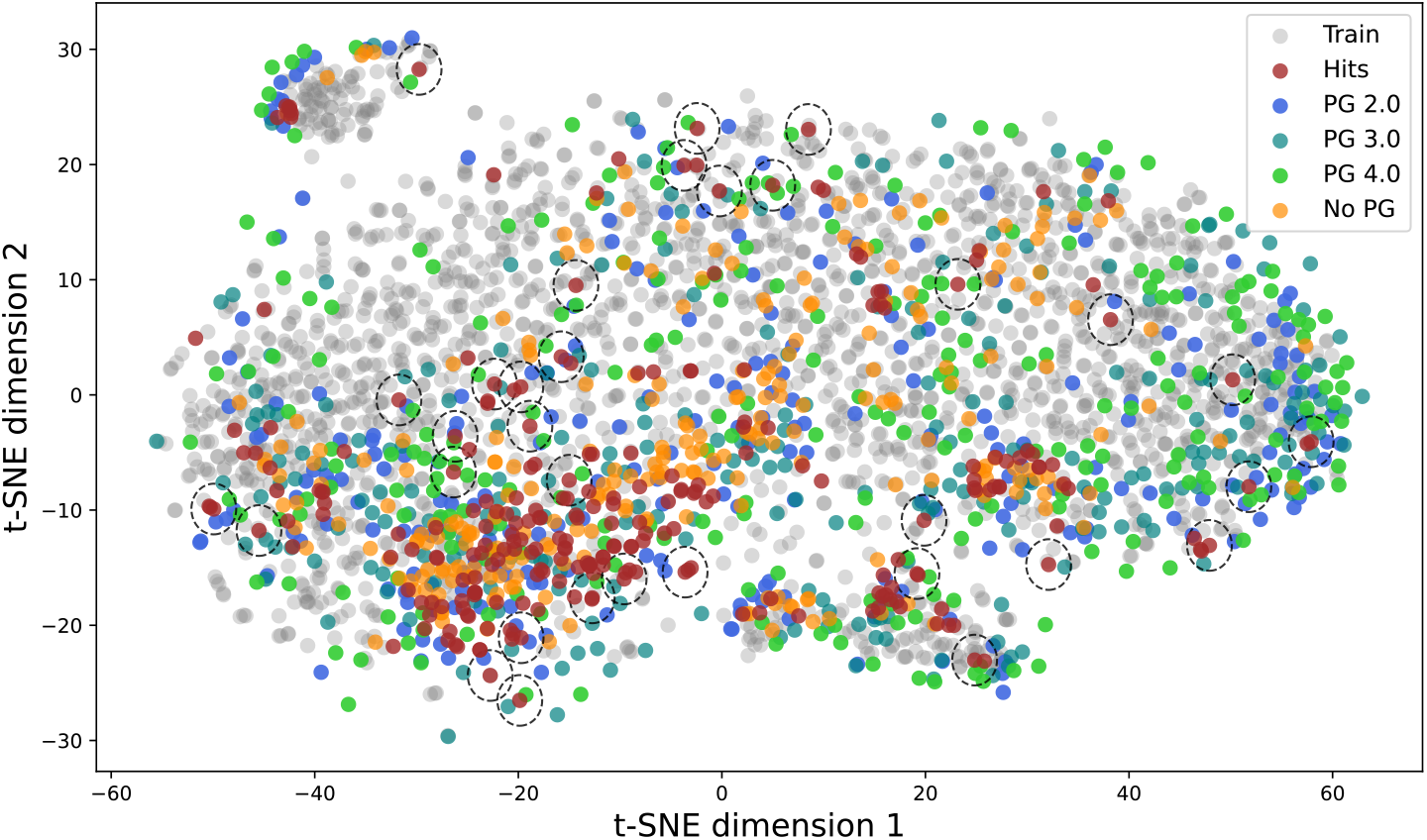
A t-SNE of dataset samples, hits, samples conditioned for potency, and samples conditioned for potency with particle guidance. Particle guidance improves sampling of the smaller hit clusters, without sacrificing focus on potency which is a useful generalization of the classic and often invoked sphere exclusion clustering technique.

## 6 Conclusions

In this work, we present COATI-LDM, a novel latent diffusion framework for small molecule design and optimization. We show that latent diffusion can be applied directly and conveniently using conditioned diffusion, classifier-free guidance, and classifier guidance model schemes. We demonstrate that such schemes can accurately model molecular data distributions and provide control over molecular properties of interest, such as physicochemical properties and binding affinities, with competitive quality to available alternatives. Furthermore, we demonstrate the practical utility of this framework, such as the ability to simultaneously generate close analogs that improve upon molecular properties and diverse generation of chemical sets using a particle guidance approach. Additionally, we utilize an open-source medicinal chemistry preference dataset to align generations to medicinal chemists’ interests and validate these generations using a prospective survey of trained chemists. In sum, this latent diffusion methodology offers a practical and effective tool for conditional molecular generation, with clear applications in the hit-to-lead and lead optimization stages of small molecule drug discovery.

## A Human Carbonic Anhydrase II Data

The hCAII binding affinity data is produced using our in-house, proprietary tArray platform, which measures quantitative binding affinities of millions of small molecules to a biological target of interest (protein, RNA, etc.) through an image-based, fluorescent readout. The small molecule libraries are built combinatorially using commercial and custom building blocks - with the small molecules tethered to beads that are immobilized to a 32 million-well silicon chip that is the size of a nickel. The targets are either directly labeled with a fluorophore or detected through a fluorescently-labeled secondary detection modality (i.e., Anti-6X His Tag Antibody) that binds the complementary tag on the target. Each target-molecule interaction is measured 15 times per chip on average (governed by a Poisson distribution). 500k rows of data were randomly selected from this data for our experiments. Binding is regressed as a function of the cumulative distribution function (CDF) in our experiments. This data will be made available at https://github.com/terraytherapeutics/COATI-LDM at a later date.

## B Encoder-Decoder Details

Our molecular encoder-decoder is derived from the architecture of [Kaufman et al., 2024], with a few architectural changes and an expanded training set. The E(3)-GNN architecture used for the 3D conformer encoder was replaced with an Allegro model [Musaelian et al., 2023], which provides resolution of chirality. The size of the transformer encoder was increased up to 62M parameters. We used 512-dimensional encodings for all experiments in this work (up from 256 in the previous architecture). We replace the InfoNCE [Oord et al., 2018] and Barlow Twins [Zbontar et al., 2021] losses used in the original work with DirectCLR [Jing et al., 2022], which we find empirically to reduce dimensional collapse in the latent representation.

The sources included in the encoder-decoder dataset (Table 5) were mostly synthetically reasonable catalogs (Enamine, Mcule, Wuxi, Zinc22, Terray Libraries, and Enamine building block space) but supplemented with compounds from in-vitro & in-vivo studies (Gostar, CHEMBL), conformer ensembles from GEOM-Drugs [Axelrod and Gómez-Bombarelli, 2022] & TensorMol [Yao et al., 2017], and permuted compounds (Protonation Enumeration, ChEMBL 33 CREM, Difficult CREM, Multi-Fragment) via CREM [Polishchuk, 2020] or Dimorphite-DL [Ropp et al., 2019].

**Table 5:** Sources sampled to train the encoder-decoder.

| Name | Description | Samples |
| --- | --- | --- |
| Enamine Real | Classical database of enumerated structures | 59,999,028 |
| Mcule | Purchasable Mcule supplier & ULTIMATE catalogs | 39,999,889 |
| Virtual Terray Libraries | Virtually enumerated molecules available to Terray’s screening assay | 39,999,708 |
| Wuxi | Virtual screening catalog | 39,953,341 |
| Zinc22 | Database of commercially-available compounds for virtual screening | 35,262,038 |
| Terray Libraries | Sample of combinatorial libraries used in Terray’s screening assay | 13,999,674 |
| Gostar | Curated SAR database | 9,088,018 |
| Protonation Enumeration | ChEMBL and Terray library samples modified using Dimorphite-DL | 8,147,175 |
| GEOM-Drugs | Molecules with multiple energy-annotated conformations | 6,999,873 |
| ChEMBL 33 CREM | ChEMBL molecules modified using the CREM package | 5,235,347 |
| Difficult CREM | Low COATI[Kaufman et al., 2024] likelihood molecules modified using CREM | 2,308,381 |
| ChEMBL 33 | Curated database of bioactive molecules with drug-like properties | 1,325,368 |
| TensorMol | Molecules with multiple DFT-optimized conformers | 399,946 |
| Enamine Building-Blocks | Combinatorial library building blocks from Enamine | 299,953 |
| Multi-Fragment | Two fragment molecules from ChEMBL | 58,118 |

**Table 6:**
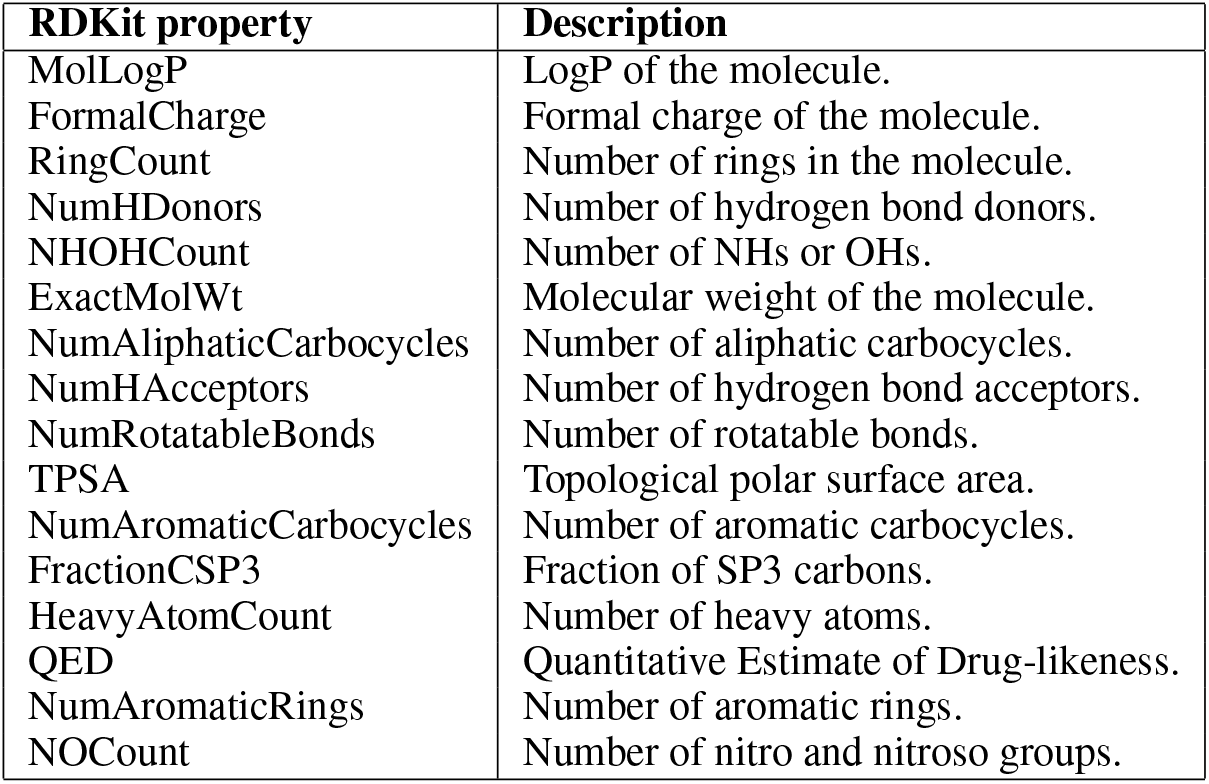
The list of specific RDKit properties included as tokens in the COATI encoder-decoder transformer model.

As described in the main text, the molecular decoder we employ can complete prompts that contain a property condition. Table 6 contains a complete list of available properties which were obtained via RDKit [RDKit, online].

## C Score Network Architecture

This work replaces the convolution layers of a U-Net with 2-layer weight normalized [Karras et al., 2023] SwiGLU [Shazeer, 2020] modules which downsample and upsample. Three downsampling stages, followed by three upsampling stages gave the best results in our experiments. Any scalar variables made available to a score network are sinusoidally encoded. Unless otherwise mentioned 1000 timesteps are used in all sampling experiments with a linear noise schedule with *β* in range [0.0001, .02].

## D Preference Modeling

To test our ability to control the “desirability” of a molecule, we first train an MLP RankNet using the methodology of [Choung et al., 2023b]. We observe in Figure 11 that these latent embeddings reproduce similar performance as the original fingerprint-based model.

**Figure 11.**
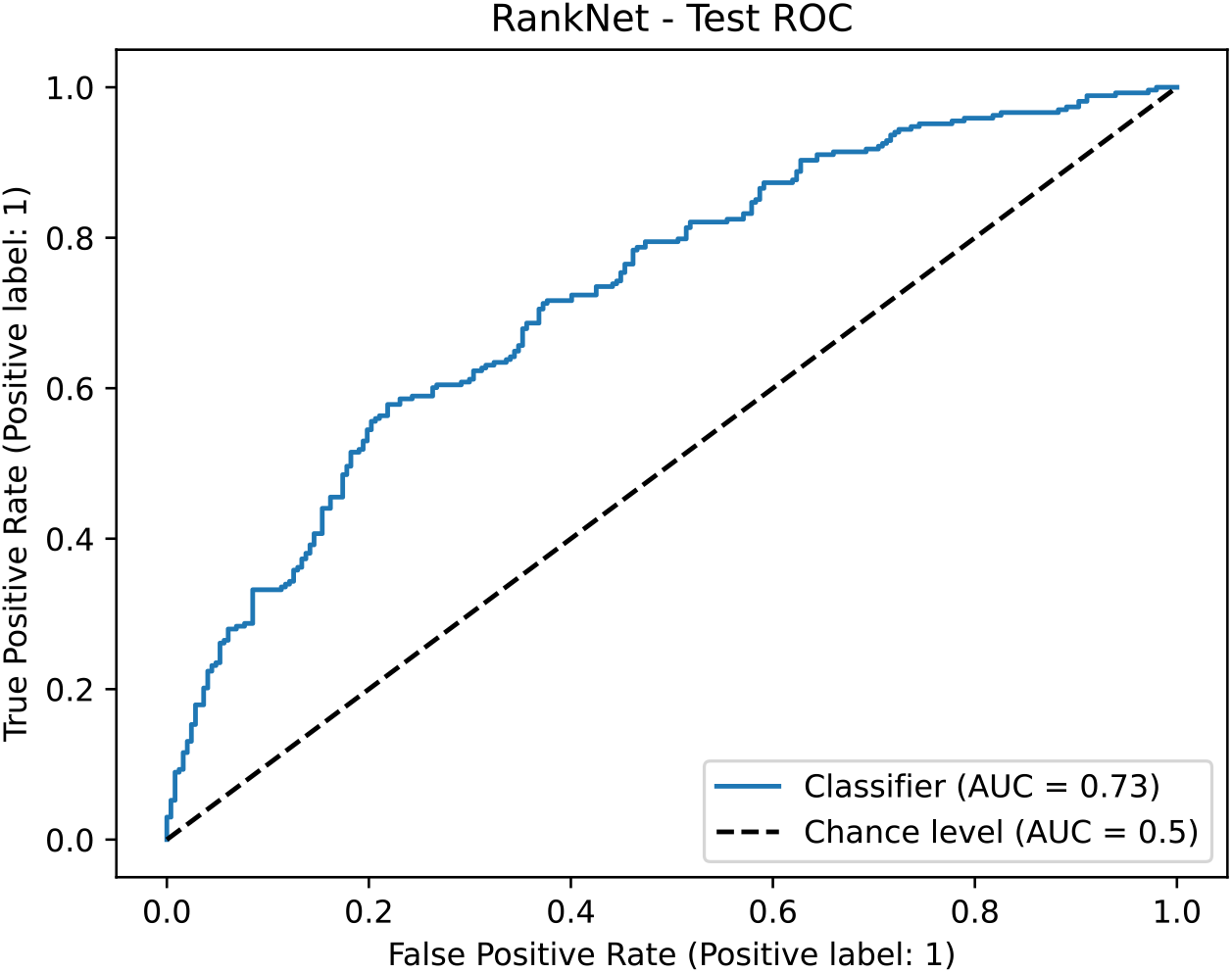
Receiver operating characteristic (ROC) curve of held-out molecule ranking pairs. This reproduces the results from [Choung et al., 2023b]

**Figure 12.**
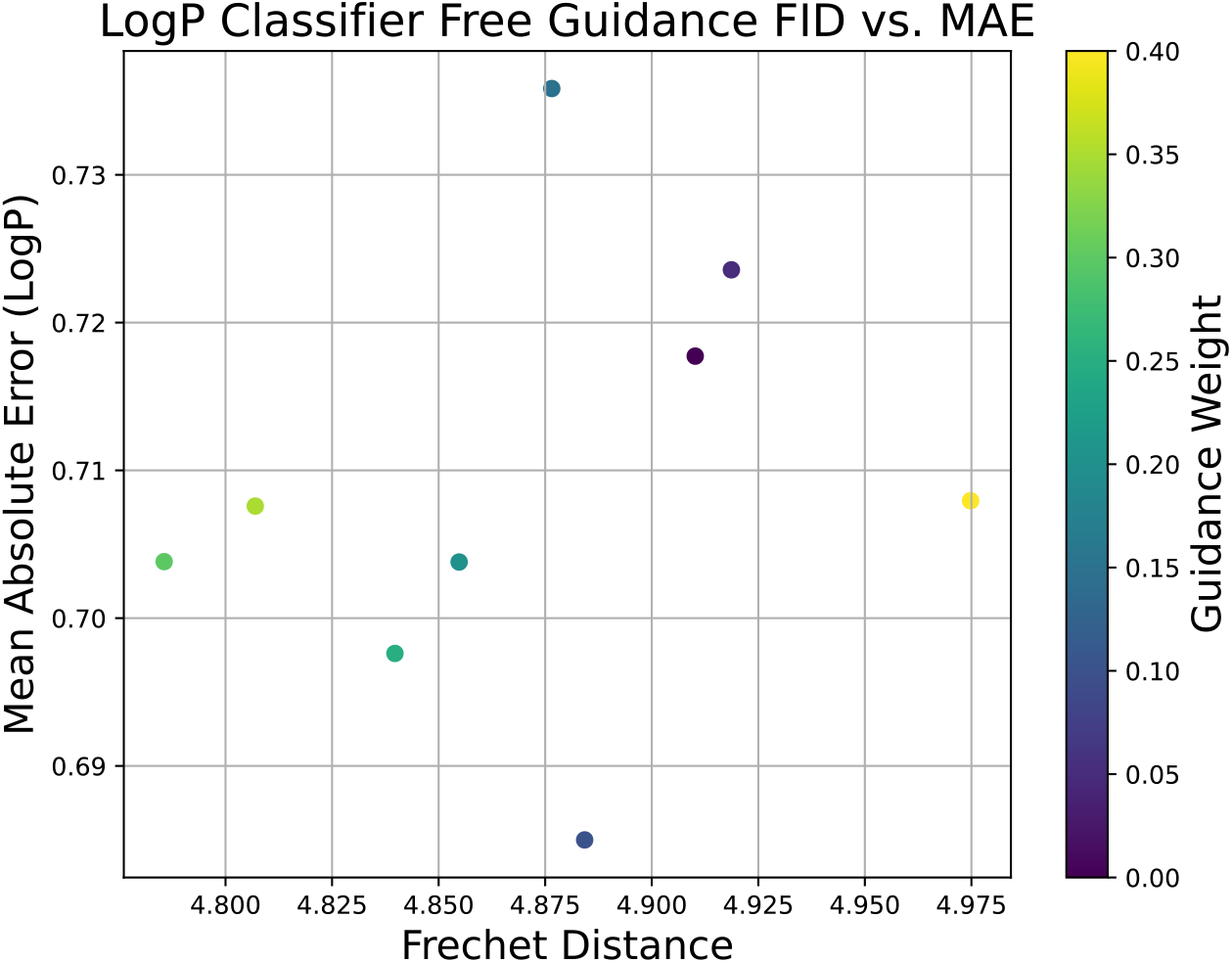
Mean Absolute Error and Frechet Distances of different Classifier-free guidance weights when conditioned on LogP. We observe no obvious correlation between FD and MAE, with a limited range of both FDs and MAEs across weights.

**Figure 13.**
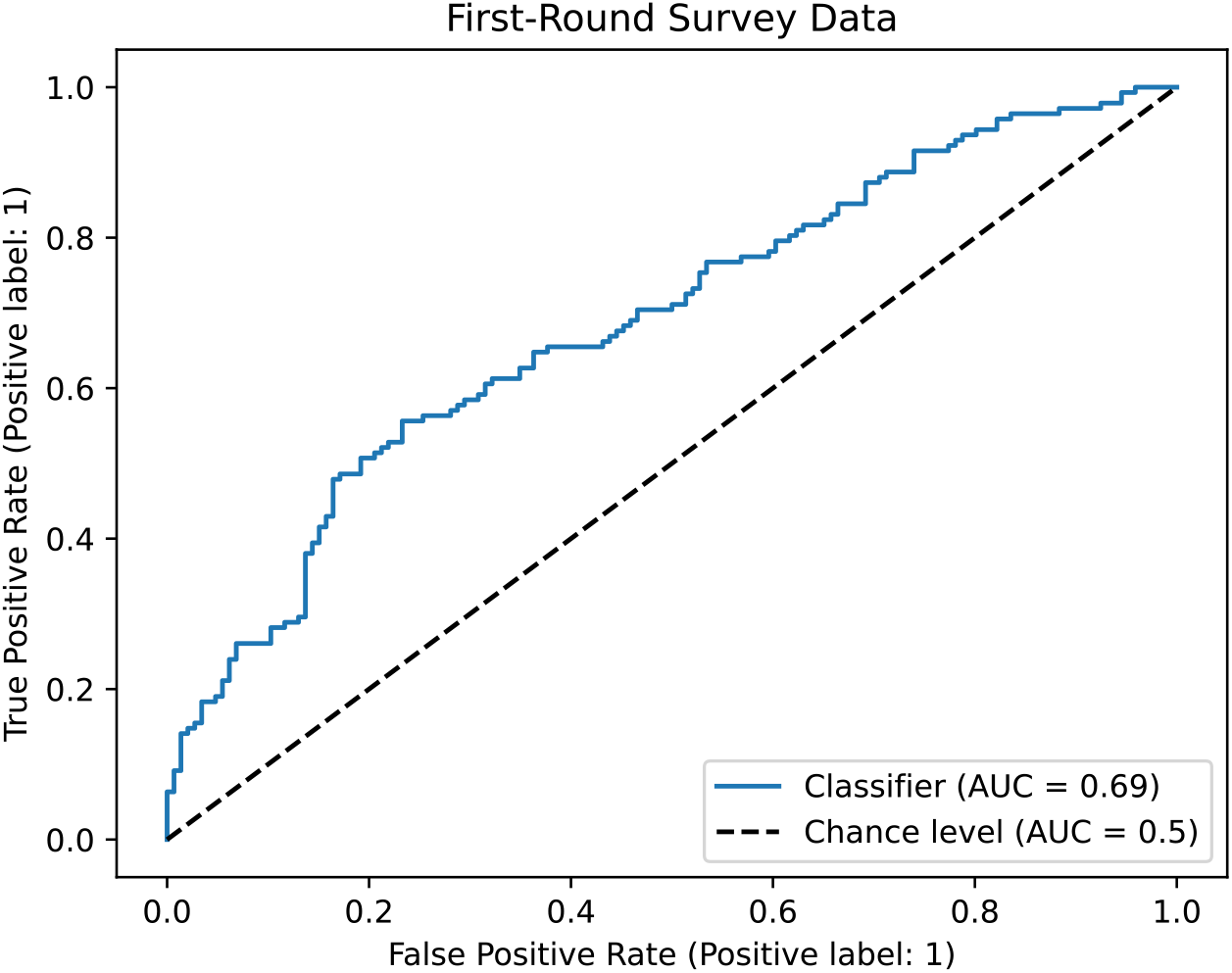
Pre-trained RankNet model inference on first round survey data. We find the preference model to have moderate agreement with our survey participants, on par with the held-out pairs from the model’s data source.

**Figure 14.**
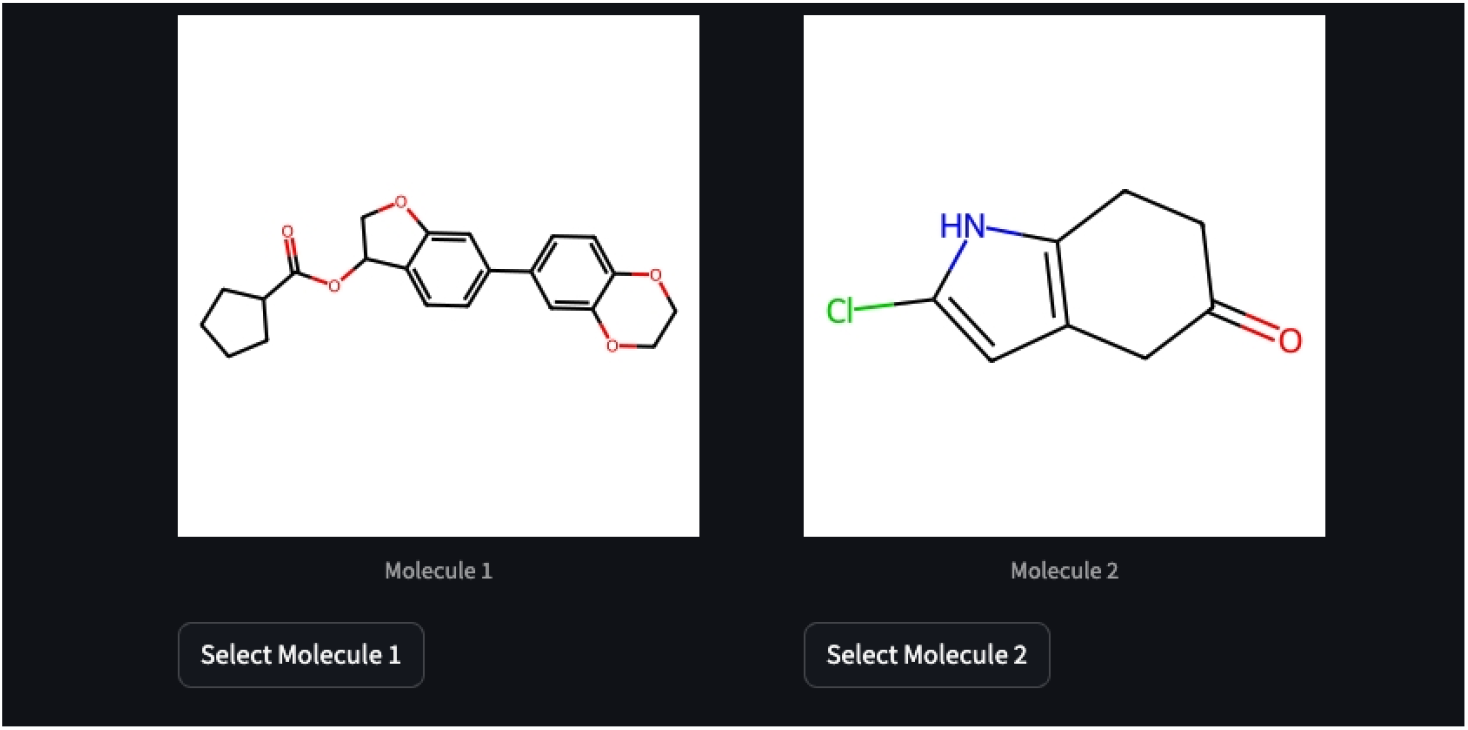
Streamlit molecule selection interface.

We can then relabel the RankNet training molecules using the model, and fit a conditional diffusion model using the procedure described in Section 3.

We see in Table 7 that control over the preference score, as measured by RankNet predictions on the resulting samples, is generally good. The conditional diffusion model can easily learn the target mapping (“copying” the RankNet scoring model) and perform conditioning of a large diffusion model trained on a very distinct dataset.

**Table 7:** Property/Target Correlations.

| Method | Preference |
| --- | --- |
| Joint | .923 |
| Classifier-Free Guidance | .938 |
| Classifier Guidance | .770 |

Given that one of the major benefits of this preference modeling is its improvability, we investigated potential deficiencies in the scoring or generation. Feedback from participants indicated that some pairs were ambiguous, and a qualitative analysis of the pairs where chemists disagreed with the model show strange or otherwise unstable groups. Figure 15 shows an example pair where the respondent chose the low-preference generation over the high-preference generation. We observe that the high-preference generation includes a highly strained ring system, indicating that the learned preference score has an incomplete understanding of chemical stability. However, this feedback can be easily integrated into the learned preference framework.

**Figure 15.**
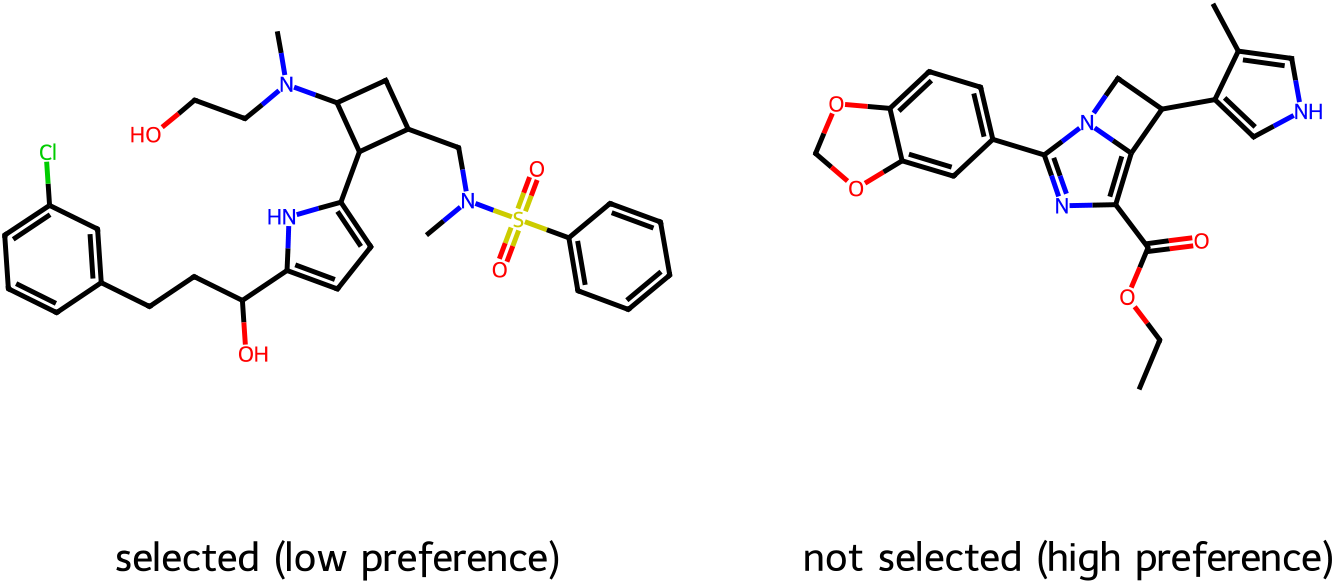
A pair of molecules where the low-preference generation was chosen. We observe that the high-preference generation includes a highly strained ring system, indicating that the learned preference score has an incomplete understanding of chemical stability.

We hypothesize that the preference model may be incompletely aligned, or conditioning using such a small training dataset could lead to degenerate samples. In aggregate, we find these proof-of-concept results encouraging, in particular due to the improvability of our preference model. We anticipate exploring the adaptation of chemical preference alignment to the needs of specific teams or projects in future work.

## E QED benchmark methodology

For the nearby diffusion on the QED benchmark task we used the same unconditional latent generator trained on the data in table 5 and trained a DUE Classifier Guide on a random 500k subset to predict QED. we embedded the 800 molecules in the benchmark set with our encoder and then applied a grid-search like methodology to each of them up to the maximum 50,000 oracle calls specified by the benchmark. The pseudocode for this approach is in algorithm 1.

### Algorithm 1

QED Benchmark Approach

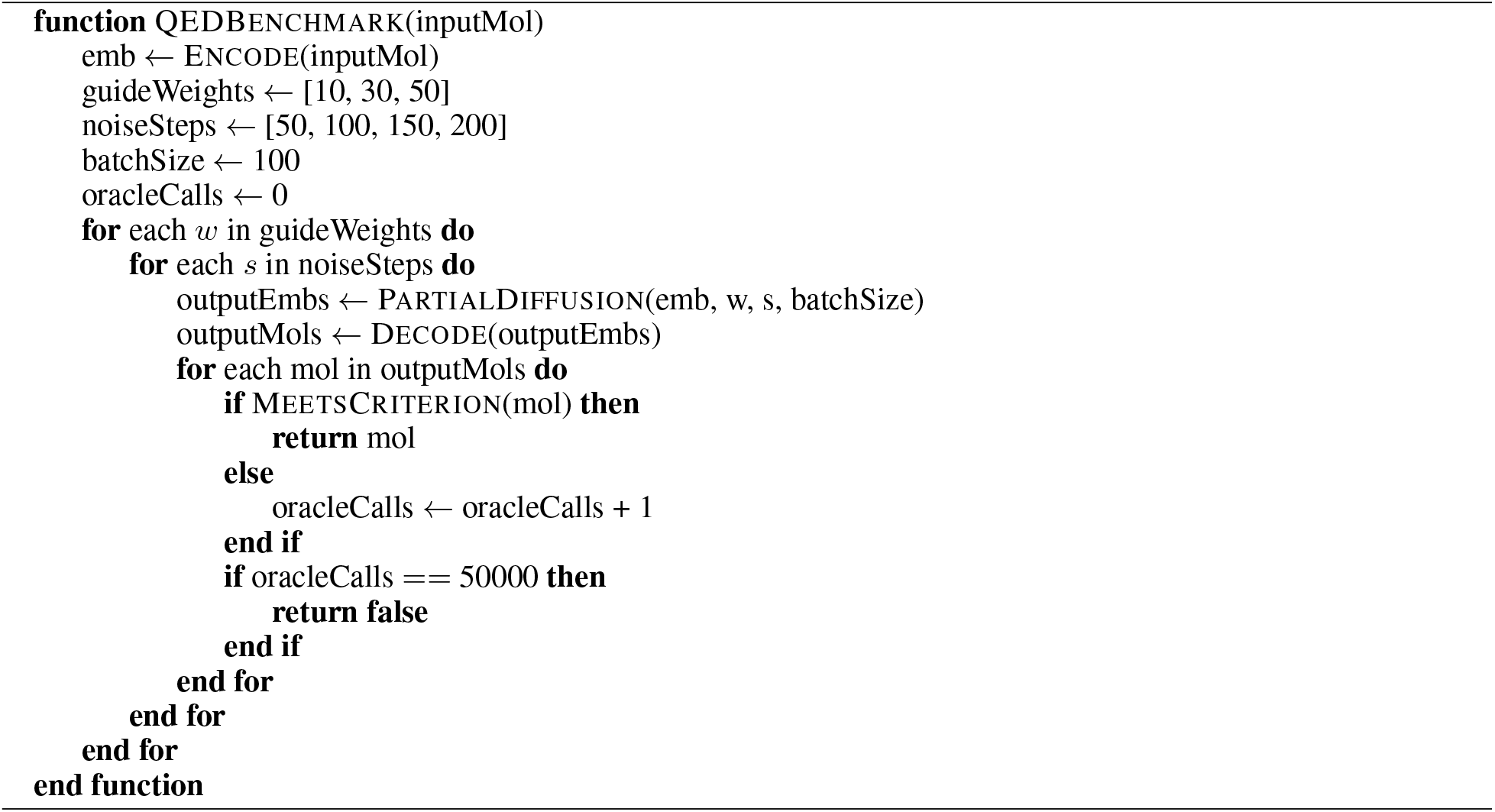

